# Predictive stability code: Trial-by-trial conflict expectations tune task representations via proactive and reactive spatio-temporo-spectral dynamics

**DOI:** 10.64898/2026.09.20.752953

**Authors:** Giada Viviani, Antonino Visalli, Maria Montefinese, Irene Di Pietro, Antonino Vallesi, Maria Ruz, Ettore Ambrosini

## Abstract

Cognitive stability, or task focus, enables goal-directed behavior through representational dynamics: by prioritizing task-relevant representations over competing task-irrelevant ones. Beyond corrective reactive mechanisms operating after conflict arises, stability can also be engaged predictively through the statistical learning of conflict regularities, either proactively, based on global conflict expectations, or through a predictive reactive mode, guided by item-specific conflict expectations. Yet, the representational mechanisms enabling such predictive adjustments remain underspecified. We addressed this by re-analyzing EEG data from a spatial Stroop task concurrently manipulating list-wide (LWPC) and item-specific (ISPC) proportion congruency. Moving beyond existing analytical approaches, we developed a framework that combines trial-by-trial Bayesian conflict-expectancy modeling with a novel spatio-temporo-spectral searchlight Representational Similarity Analysis, and leveraged directional representational modeling complemented by ridge-regression decoding. We first identified the necessary computational substrate for predictive stability reconfigurations: the spatio-temporo-spectral patterns encoding task (Target, Distractor) and conflict-expectancy (LWPC, ISPC) representations, with the latter already encoded in the pre-stimulus windows. Crucially, conflict expectations modulated Target and Distractor encoding strength with distinct spatio-temporal and spectral dynamics, revealing a functional dissociation. LWPC-induced proactive stability follows a “Target-first” policy, primarily strengthening the Target representation early after stimulus onset via Alpha and lower-Beta patterns, and weakening Distractor encoding only later; conversely, ISPC-induced predictive reactive stability follows a “Distractor-first” policy, rapidly weakening Distractor encoding via response-locked posterior Beta patterns, and strengthening the Target representation only downstream. Our study refines theoretical cognitive control models — incorporating a mechanistic explanation for predictive stability — while also establishing a multidimensional methodological template for investigating the underlying substrates of adaptive behavior and beyond.

## 1. Introduction

Human goal-directed behavior critically relies on the ability to adaptively coordinate thoughts and actions in response to changing goals and contextual demands, a capacity broadly known as cognitive (or adaptive) control (Chiew & Braver, 2017; Cohen, 2017; Miller & Cohen, 2001). A core component of this ability is cognitive stability (or task focus), as opposed to flexibility (or switch readiness), which enables the system to handle conflict by prioritizing currently relevant tasks over competing, irrelevant information or tasks (Egner, 2023). Traditionally, stability has been conceptualized as a corrective reactive mode —a transient mechanism engaged “just-in-time” to resolve interference after conflict is detected (Braver, 2012; Braver et al., 2007). However, stability is an adaptive capacity that does not solely rely on late corrections; rather, it also recruits predictive mechanisms that dynamically adjust the system to match contextual demands through the statistical learning of contextual regularities, such as the probability of conflict. These predictive stability mechanisms operate upstream to preemptively minimize the impact of conflict (Egner, 2023; Jiang et al., 2014). Predictive regulation can be implemented through a proactive stability mode, as formalized by the Dual-Mechanisms of Control (DMC) model (Braver, 2012), which distinguishes between proactive and reactive modes of control modes^1^. Specifically, proactive stability anticipates conflict through the sustained upregulation of task focus when high global conflict is expected. Furthermore, as elaborated in our recent work (Viviani et al., 2026), stability can also manifest in a predictive reactive mode: this mode is triggered by stimulus presentation but remains fundamentally predictive, as it uses learned item-specific conflict expectations to strategically prepare for conflict even before it fully emerges. To clarify terminology, we refer to the reactive mode originally proposed by the DMC as corrective reactive stability, distinguishing it from the predictive counterpart mode. Together, these three modes highlight the remarkable adaptability of stability across different contexts and timescales. In this study, we use multivariate analytical approaches to uncover the electrophysiological dynamics enabling the implementation of the two predictive stability modes.

Despite extensive research, the neural and computational implementation of predictive stability—how the brain preemptively sustains and adapts task focus to contextual demands— remains poorly understood (Egner, 2023; Jiang et al., 2014). A critical limitation of traditional measurement approaches is that they often infer stability implementation from its downstream effects, such as improved performance, reduced conflict, or changes in neural activation, without directly characterizing the computational substrates implementing these effects (Freund et al., 2021a). In our recent EEG study (Viviani et al., 2026), we began to address this gap by concurrently manipulating proactive and predictive reactive stability to map their high-resolution temporal (ERP) and spectral (ERSP) dynamics. Our findings showed that successful engagement of predictive mechanisms —specifically proactive stability— systematically reduced post-stimulus conflict-related activity, suggesting a diminished need for corrective reactive stability (in line with Banich, 2009). However, while these results suggest that upstream predictive regulation can pre-emptively bypass the need for costly downstream corrective processes, they remain agnostic to the neural representations the system operates on to achieve such outcomes.

A promising way to address this gap is to explicitly consider neural representations, as stability is enacted by modulating these representations. Neural representations are the symbolic neural counterpart of the information required for successful performance (Baker et al., 2022), bridging brain activity and the cognition underlying goal-directed behavior (Braver, 2012; Diedrichsen & Kriegeskorte, 2017). These representations consist of distributed multivariate patterns of neural activity that encode and transform informational content necessary to coordinate and modulate downstream perceptual, motor, and cognitive processes (Badre et al., 2021; deCharms & Zador, 2000; Diedrichsen & Kriegeskorte, 2017; Freund et al., 2021b; Kriegeskorte & Diedrichsen, 2019; Kriegeskorte & Kievit, 2013; Sakai, 2008). Both theoretical accounts and empirical evidence suggest that stability implementation relies on neural codes encoding task and conflict-expectancy representations (e.g., Badre et al., 2021; Cellier et al., 2022; Jiang et al., 2018). Specifically, stability operates by actively maintaining task (or task set) representations —neural patterns encoding the goals and rules that link stimuli to appropriate responses (Kiesel et al., 2010; Monsell, 2003). Stability is thus assumed to be implemented by selectively enhancing the encoding strength of the relevant task representation, shielding it from interference by competing, irrelevant task representation(s). This representational bias is then used by downstream processes to prioritize task-relevant information and facilitate correct response selection (Cellier et al., 2022; D’Esposito, 2007; Egner, 2023; Miller & Cohen, 2001; Nee et al., 2007; Sakai, 2008; Schumacher & Hazeltine, 2016). However, while this representational account is increasingly implicit in major theoretical frameworks of cognitive stability (e.g., Braver, 2012; Egner, 2023; Miller & Cohen, 2001), these frameworks were not originally formalized in representational terms —that is, they do not specify the neural codes they invoke and the corresponding encoding strength and its directional modulations. Empirical literature has primarily measured correlates of stability-related processes through aggregate neural signals rather than through explicit representational modeling. This leaves a gap between the theoretical postulation of representational mechanisms and their direct empirical characterization.

The different modes of stability may target these task representations at different stages. Corrective reactive stability intervenes as a late mechanism to resolve occurring conflict, likely through just-in-time reactivation of relevant task representations and downstream motor adjustments, such as selecting the correct response over competing alternatives during the final stages of the selection cascade (Banich, 2009; Viviani et al., 2026). In contrast, predictive stability mechanisms are expected to operate primarily upstream, relying on the encoding of conflict-expectancy representations —internal estimates of conflict probability derived from experience (e.g., Egner, 2023; Jiang et al., 2018). By determining the current stability demand, these expectations are continuously updated and used to predictively modulate the encoding strength of the task representations, either proactively or reactively (e.g., Cellier et al., 2022; D’Esposito, 2007; Egner, 2023; Jiang et al., 2018). Therefore, to understand stability, especially in its predictive forms, it is essential to focus on how conflict-expectancy representations modulate the neural code strength of task representations before downstream correction becomes necessary.

As noted above, despite their central theoretical role, these representational dynamics have rarely been directly investigated in the study of cognitive stability, which has likely contributed to the lack of a comprehensive mechanistic account after half a century of research on cognitive control (Freund et al., 2021a). A key reason is that most previous research —including our own prior work (Viviani et al., 2026)— has relied on traditional mass-univariate analytical approaches. These methods aggregate neural activity across a small number of abstract factor levels (i.e., experimental conditions), indexing only the overall magnitude of neural activation and its macroscopic changes across experimental manipulations. While this approach can reveal the extent to, and conditions under which stability-related processes are engaged (Cheng, 2021) —their downstream consequences— it discards the rich informational content encoded in distributed neural activity patterns —the candidate upstream computational mechanisms (Diedrichsen & Kriegeskorte, 2017; Kriegeskorte & Diedrichsen, 2019; Popal et al., 2019). To overcome these limitations, it is therefore necessary to move beyond measuring aggregate processes toward explicitly modeling neural representations using multivariate pattern analytical techniques.

### 1.1. Multivariate pattern analysis of stability representations

Multivariate pattern analysis (MVPA) offers a robust framework to investigate the representational mechanisms underlying stability by interrogating the high-dimensional neural code embedded in distributed activity patterns (Diedrichsen & Kriegeskorte, 2017; Kriegeskorte & Kievit, 2013). While initially developed for fMRI, MVPA is particularly powerful when applied to M/EEG data, as it harnesses high temporal resolution to capture the inherent dynamic nature and interactions of representations over time (Cellier et al., 2022; Fahrenfort et al., 2017, 2018).

Two complementary families of MVPA methods exist: decoding and encoding. Decoding models use neural activity patterns as input to predict informational content (e.g., experimental conditions, stimulus features). The successful decodability of a variable suggests that the relevant information is contained and accessible within those multivariate patterns (Kriegeskorte & Diedrichsen, 2019; Popal et al., 2019). Traditionally, decoding has been achieved through linear classifiers, which identify the discriminability between discrete categories (e.g., Duda et al., 2001; Mahmoudi et al., 2012). However, classification-based approaches are often limited by their categorical nature, which can restrict the exploration of more complex, continuous informational spaces. Regression-based decoding, including regularized methods such as ridge regression, addresses this limitation by enabling fine-grained predictions of continuous variables (Bode et al., 2021; Cohen et al., 2011).

Despite its widespread employment, decoding relies on reverse mapping, which can leave the researcher blind to the specific information driving decodability, thus preventing a full characterization of the multidimensional structure (i.e., representational geometry) of neural representations (Diedrichsen & Kriegeskorte, 2017; Kriegeskorte & Diedrichsen, 2019; Ritchie et al., 2019). These limitations are addressed by encoding models, which adopt a forward inference approach, using descriptions of the informational content (experimental conditions/stimuli) as input to predict brain patterns (Kriegeskorte & Douglas, 2019). The most common of these, Representational Similarity Analysis (RSA), explicitly models the representational geometry of theoretically defined models and tests them against the observed neural similarity structure (Kriegeskorte et al., 2008). By quantifying the (dis)similarity between the representational geometries of neural patterns and predicted models through Representational Dissimilarity Matrices (RDMs), RSA allows for a complex characterization of the representational space, enabling the quantification of the encoding strength with which specific theoretical elements —such as task or conflict-expectancy representations— are represented in the brain (Diedrichsen & Kriegeskorte, 2017; Freund et al., 2021a; Kriegeskorte & Kievit, 2013).

In recent years, multivariate techniques have begun investigating the representational architecture of cognitive stability, revealing its reliance on the active maintenance and reconfiguration of specific multivariate neural codes. Early fMRI classification-based decoding studies demonstrated that frontoparietal networks maintain and reconfigure abstract representations of task rules (Qiao et al., 2017; Woolgar et al., 2011) and encode conflict as a distinct, generalizable representation (Jiang & Egner, 2014). More recently, RSA has been used to map the representational geometry of stability-related information. For instance, applying RSA to fMRI data within a color-word Stroop task, Freund and colleagues (2021b) investigated the neural representations of target (color), distractor (word), and conflict information. They showed that the representation of goal-relevant information (the target) in the dorsolateral prefrontal cortex was associated with reduced Stroop interference, consistent with a role in proactive stability, and that this operated alongside a distinct representation of conflict in the dorsomedial frontal cortex. Furthermore, conflict representations have been found to be organized within a structured cognitive space related to the magnitude of trial-by-trial behavioral adaptations (Yang et al., 2024). Complementing these spatial insights, recent EEG studies have employed RSA to track the temporal dynamics of task representations (or conjunctive representations) that bind stimuli, rules, and responses, showing that these have a hierarchical structure and are stably maintained over time to continuously bias behavior (Cellier et al., 2022; Kikumoto et al., 2022a). More recently, Gheza et al. (2026) applied a decoding-RSA approach to test how target and distractor representations are modulated by conflict history at two temporal scales: trial-by-trial (based on the previous trial’s congruency) and long-term (through a task-specific proportion congruency manipulation). They found reactive suppression of the distractor —but no target modulation— with long-term learning tuning the efficiency of this stimulus-triggered suppression rather than establishing anticipatory representational states.

Together, this emerging literature confirms that stability operates by modulating specific multivariate patterns encoding information about the task and conflict. However, a comprehensive mechanistic account of how predictive stability dynamically and anticipatorily reconfigures neural representations to prevent conflict has remained elusive. Due to several theoretical and methodological constraints, existing approaches have neither directly tested the directional modulation of task representations by continuously updated conflict expectations, nor characterized the full spatio-temporo-spectral dynamics through which this reconfiguration unfolds —leaving the representational implementation of predictive stability largely underspecified.

#### 1.1.1. Theoretical limitations

MVPA research on cognitive stability has primarily focused on corrective reactive stability. Most studies have revealed either the brain’s representation of conflict as a post-stimulus outcome (e.g., Jiang & Egner, 2014; Yang et al., 2024) or task representations in isolation (e.g., Woolgar et al., 2011), largely ignoring the upstream proactive and predictive reactive mechanisms that allow the system to mitigate conflict before it emerges, adapting to contextual demands. Even when experimental designs allow for the investigation of both predictive stability modes (e.g., the DMCC project, which manipulated both global and item-specific conflict expectancy; Braver et al., 2021), existing MVPA analyses often focus solely on proactive stability, neglecting its predictive reactive counterpart (Freund et al., 2021b; see also Gheza et al., 2026, who used a different experimental paradigm). Consequently, it remains unknown how different sources of conflict expectancy interact to shape the neural code of task representations.

Second, existing research often relies on an assumption of static stability, neglecting its inherently adaptive nature. Studies typically treat stability demands as fixed states tied to experimental blocks (e.g., Freund et al., 2021b), assuming that participants immediately switch stability demand as conditions change. This “on/off” switch logic ignores that real-world stability relies on continuous statistical learning, where conflict expectations are updated trial-by-trial (Di Pietro & Viviani, 2026; Jiang et al., 2014, see also Viviani et al., 2026). While recent studies have successfully tracked task representations at the single-trial level (Cellier et al., 2022; Kikumoto et al., 2022a), they have focused on the “what” (task features) without modeling the “why” (the underlying conflict expectations). A partial exception is Gheza et al. (2026), who tested how prior conflict —indexed either by the immediately preceding trial or by long-term block statistics— modulates task representations; however, both indices treat expectations as discrete snapshots/summaries of conflict history rather than as continuously updated internal estimates. By failing to dynamically model the fluctuations in conflict expectancy, these approaches cannot identify the proximal drivers of task representational reconfiguration.

Therefore, to the best of our knowledge, no study has yet adopted an integrated approach for proactive and predictive reactive stability, investigating how conflict-expectancy representations — maturing trial after trial— sculpt the neural code of task representations before conflict occurs.

#### 1.1.2. Methodological limitations

Addressing the aforementioned theoretical gaps requires overcoming several methodological constraints that have limited previous investigations.

First, while the inherent limitations of decoding described above have been overcome by RSA studies, these have predominantly relied on fMRI (e.g., Freund et al., 2021b; Yang et al., 2024), which lacks the sub-second temporal resolution necessary to reveal the temporal trajectories of dynamic representational reconfigurations. Consequently, it does not allow for capturing the continuous, trial-by-trial updating of conflict expectations that drive stability adaptations, nor for tracking whether and how these expectations lead to the anticipatory (proactive stability) or stimulus-triggered (predictive reactive stability) modulation of task representations.

Second, even when the high-temporal resolution of EEG is leveraged, the multivariate signal is typically collapsed across dimensions. Studies commonly adopt a time-resolved approach —where brain RDMs are computed for each time point using the whole spatial pattern of EEG activity (e.g., Kikumoto et al., 2022b; Gheza et al., 2026). By relying on global spatial patterns —which has inherently low resolution in EEG— as the sole multivariate feature to draw inferences about temporal dynamics, this method remains blind to topographical specificity. Crucially, this single-feature approach also fails to exploit the temporal patterns of the neural signal, effectively discarding the fine-grained temporal information that constitutes EEG’s primary strength. Moreover, it prevents determining how spatial and temporal features jointly encode stability-related representations. A multi-feature searchlight approach integrating space and time (Higgins et al., 2022) can overcome this issue by using spatial and temporal information simultaneously, allowing for both spatial and temporal inference; however, it has not yet been systematically applied to cognitive stability.

Third, beyond space and time, the spectral domain is systematically overlooked, despite neural oscillations carrying rich, content-specific representational information (Kikumoto & Mayr, 2018). Failing to integrate spectral patterns into multivariate models thus means discarding a fundamental dimension of neural representational dynamics (Sommer et al., 2022). Cellier and colleagues (2022) used a time-frequency resolved RSA. However, while their analysis was resolved across frequencies, it relied on the entire spatial topography as the sole multivariate feature within each time-frequency bin. Consequently, this approach neglected the representational content potentially carried by the spectral patterns themselves and remained blind to the spatial patterns encoding representations. Therefore, to date, no study has yet employed a complete multi-feature searchlight approach — simultaneously integrating space, time, and frequency— to obtain a fully multidimensional picture of stability representational dynamics.

Fourth, existing multivariate approaches typically quantify how control shapes task representations in terms of average encoding strength across trials, which reveals whether a representation is modulated but not the direction of the modulation. To determine whether higher conflict expectations strengthen the target and weaken the distractor (or vice versa), it is necessary to use analytical tools that explicitly rank trial pairs along the expectation scale, an approach that, to our knowledge, has not yet been implemented.

Lastly, an opportunity for further methodological integration lies in combining representational similarity analysis (RSA) and decoding approaches. While RSA can characterize the geometry of representational spaces and decoding can assess the information carried by neural patterns, integrating these complementary methods could provide converging evidence and offer a more comprehensive characterization of the identified representations.

### 1.2. The present study

The present study provides a multivariate re-analysis of the EEG dataset previously reported in Viviani et al. (2026). In that work, mass-univariate analyses (ERPs and ERSPs) revealed a robust effect of proactive stability which, when highly engaged, systematically reduced neural markers of conflict detection and resolution —reflecting a diminished need for a late-stage, just-in-time braking mechanism (i.e., corrective reactive stability; see Tafuro et al., 2020). While we failed to observe such an effect for predictive reactive stability, our previous behavioral evidence (from a larger sample, see Viviani et al., 2024a) has shown that it comes into play when proactive stability is less engaged. As we speculated in Viviani et al., 2026, the failure to find clear univariate effects for this mode could stem from the fact that it operates primarily at the representational level, rather than through large-scale neural processes. The apparent discrepancy between univariate null findings and multivariate sensitivity for predictive reactive stability reflects a fundamental difference between these analytical approaches: mass-univariate measures (ERPs and ERSPs) capture broad, net changes in population-level neural synchronization —well-suited for global, sustained shifts such as those induced by proactive stability— but insensitive to the fine-grained spatial and spectral reconfigurations hypothesized to underlie item-specific regulation, which might instead selectively modulate distinct neural sub-populations without altering overall macroscopic power. Overall, while our previous study provided crucial insights into the downstream consequences of predictive stability, it could not reveal the upstream neural dynamics of strategic conflict mitigation. Therefore, our core objective here is to provide a comprehensive mechanistic account of how the brain implements both predictive stability modes by shifting the focus directly onto the representational dynamics that allow the system to anticipatorily configure the neural code of the task.

To achieve this, here we investigate the adaptive nature of predictive stability at two levels. First, we examine how the cognitive system adaptively deploys different predictive stability modes — proactive or predictive reactive— based on the specific conflict probabilities available in the environment. To test this, the perifoveal spatial Stroop paradigm used in Viviani et al. (2026, and validated in Viviani et al., 2024a) —in which participants identified the direction of a target arrow while ignoring its distracting spatial position— was ideal for inducing both modes simultaneously to investigate how these two predictive mechanisms coexist to meet different stability demands. Within it, global and item-specific probabilities of conflict were concurrently manipulated to induce distinct conflict expectancies, thereby varying proactive and predictive reactive stability demands. Specifically, proactive stability was induced via a list-wide proportion congruency (LWPC), where global conflict probability (i.e., in the block) encourages a sustained upregulation of task focus. Concurrently, predictive reactive stability was induced via an item-specific proportion congruency (ISPC), where conflict likelihood is tied to specific stimulus features, allowing the system to trigger predictive adjustments upon stimulus identification. Second, we investigate adaptability as a continuous, trial-by-trial dynamic, moving beyond the implausible assumption that stability is a static, discrete “on/off” condition (Di Pietro & Viviani, 2026; Viviani et al., 2024a). Since predictive stability in the real world is dynamically adjusted via statistical learning —which continuously updates conflict expectations based on recent experience— we use an ideal Bayesian observer to model trial-by-trial updates of both global (LWPC) and item-specific (ISPC) conflict expectations. The resulting trial-level estimates serve as reliable proxies for proactive and predictive reactive stability demands, driving dynamic neural code modulations.

To uncover the upstream representational dynamics underlying predictive stability, overcoming the constraints of previous research, we leveraged the full high-dimensional nature of the EEG signal through an integrated multi-feature spatio-temporo-spectral searchlight approach. Unlike one-dimensional analyses that collapse the signal, our 3D searchlight simultaneously integrates spatial, temporal, and spectral dimensions (in the 4-30 Hz range). This approach ensures that neural oscillations are treated as rich sources of representational content, allowing the neural encoding of stability representations to be characterized by its precise timing, topographical distribution, and specific oscillatory profile in a single, integrated manner. We note that integrating the spectral domain serves a dual methodological and interpretive purpose: rather than exploring an unconstrained, high-dimensional frequency space, it provides a principled physiological anchor to map abstract representational dynamics onto canonical neurocomputational frameworks. Within stability dynamics, Theta rhythms (4–7 Hz) provide an established mechanism for signaling and tracking conflict, and coordinating trial-history integration (Cavanagh & Frank, 2014; Sauseng et al., 2005), whereas Alpha oscillations (8–12 Hz) implement functional gating by inhibition, selectively routing sensory information and shielding prioritized representations against incoming interference (Jensen & Mazaheri, 2010; Klimesch, 2012). Crucially, the Beta band (13–30 Hz) has emerged as the core oscillatory infrastructure supporting cognitive stability itself —mediating the endogenous, top-down maintenance of task goals, rule representations, and resistance to distraction (Engel & Fries, 2010; Miller et al., 2018; Spitzer & Haegens, 2017). Given that cognitive stability spans from the retention of environmental regularities to the execution of goal-directed behavior, tracking representational dynamics across this oscillatory range allows us to test how the brain dynamically orchestrates these spectral channels to implement proactive and predictive reactive stability.

This 3D searchlight approach was implemented using both Representational Similarity Analysis (RSA) and ridge-regression decoding, with an intentional asymmetry between the two. RSA was our primary analytical tool: by explicitly modelling the geometry of theoretically defined representations, we mapped and quantified the encoding strength of stability representations and, crucially, directionally characterized the task representation modulation by conflict expectations. Ridge-regression decoding was then employed as a convergent validation, testing whether the same informational content is structured in a format that could actually be “read out” by downstream cognitive processes. Accordingly, we report the RSA results, and use a descriptive conjunction with the decoding maps to identify the patterns for which the RSA results are confirmed by decoding ones, thereby providing the most robust evidence of stability representations.

Collectively, the combination of our experimental paradigm and multi-feature MVPA tools provided the necessary framework to test the mechanistic hypothesis of how predictive stability is implemented, thus explaining its effects observed in our previous mass-univariate study (Viviani et al., 2026). To this end, we explicitly model the two core classes of neural representations that act as the main actors enabling this: task representations, encompassing the relevant task (Target; arrow direction) and the irrelevant task (Distractor; arrow position); and conflict-expectancy representations reflecting trial-level expectations of conflict derived from global (LWPC) and item-specific (ISPC) conflict probabilities. By leveraging MVPA, we directly test the interplay between these two types of representations to track the dynamics subserving predictive stability modes.

#### 1.2.1. Hypotheses

Our main hypothesis is that predictive stability operates by dynamically reconfiguring the neural code strength of task representations based on current conflict expectations (e.g., by modulating their encoding strength according to the expected trial-level conflict). Therefore, we predict that, as a pre-requisite for successful performance, the Target is actively encoded. Moreover, we expect that the brain, through continuous statistical learning, encodes and trial-by-trial updates conflict-expectancy representations, already before stimulus onset, serving as informational content ready to preemptively sculpt task representation strength. As a result, higher conflict expectations should enhance the encoding strength of the Target representation and/or weaken the representation of the Distractor, thereby shielding the cognitive system from conflict before it occurs. While high proactive demand has been shown to modulate both target and distractor representational strength (e.g., Freund et al., 2021b) and to reduce distractor representational strength (Gheza et al., 2026), these effects have been documented as average differences in encoding strength across trials and have left two central aspects untested. First, the direction of the modulation (i.e., target strengthening and distractor weakening, or the opposite) has never been directly assessed concurrently in both directions. Second, conflict expectations have typically been treated as block-level states or discrete summaries of prior conflict rather than as continuously updated estimates. Our approach tests this interaction explicitly, at the trial level, and in a form that establishes how conflict expectations differentially modulate task codes (see Section 2.4.3).

Furthermore, we expect to observe for the first time a temporal dissociation between the two predictive modes: proactive stability (based on LWPC) should drive task representational adjustments available from the earliest stages of stimulus processing (and even earlier than stimulus onset), whereas predictive reactive stability (based on ISPC) should trigger rapid reconfigurations only upon stimulus identification. Given our integrated searchlight approach, these temporal predictions will be complemented by spatial and spectral insights, revealing the specific oscillatory profiles and cortical distributions that enable such representational dynamics.

## 2. Methods

The present study is a multivariate re-analysis of the EEG dataset previously reported in Viviani et al. (2026). Therefore, the following sections focus on the specific multivariate analytical pipeline, while providing a concise summary of the experimental procedures detailed in our prior work.

### 2.1. Participants

The sample consisted of 40 healthy volunteers (28 females, mean age = 24.71 years, *SD* = 3.76). All participants had normal or corrected-to-normal vision and reported no neurological or psychiatric disorders. Informed consent was obtained from all participants prior to the experiment. Further details on participants’ characteristics, sample size determination, and exclusion criteria are reported in Viviani et al. (2026). Briefly, sample size was determined via a priori power analysis targeting the primary univariate effects of interest. Nonetheless, a sensitivity power analysis revealed that it is sufficient to detect a minimum effect of d = 0.40 at a one-tailed one-sample t-test on the by-participants encoding strength values with a power = .80. Moreover, our sample size is consistent with those employed in comparable EEG-based RSA studies (e.g., Cellier et al., 2022; Kikumoto et al., 2022b).

### 2.2. Experimental task and procedure

Participants performed a perifoveal spatial Stroop task (Fig. 1) implemented in Psychtoolbox (Brainard, 1997) on Matlab (Version 2017b; The MathWorks, Inc. Natick, MA). For a comprehensive description of the paradigm and experimental manipulations, we refer the reader to Viviani et al. (2026). Briefly, participants were required to identify the direction of a target arrow (pointing up-left, up-right, down-right, or down-left) while ignoring its spatial position on the screen (i.e., the arrow could appear in the up-left, up-right, down-right, or down-left small quadrant around a central cross). The match or mismatch between the task-relevant feature (direction) and the task-irrelevant feature (position) resulted in congruent and incongruent trials, respectively. The Stroop effect (incongruent-congruent) yielded by this paradigm is large, robust and reliable, while ocular artifacts during EEG recording are minimized due to the perifoveal spatial arrangement of the stimuli (see Viviani et al., 2024b). Manual responses were provided using four keys compatible with the four arrow directions/positions (E, O, K, and D, using the left middle, right middle, right index and left index fingers, respectively).

**Figure 1.**
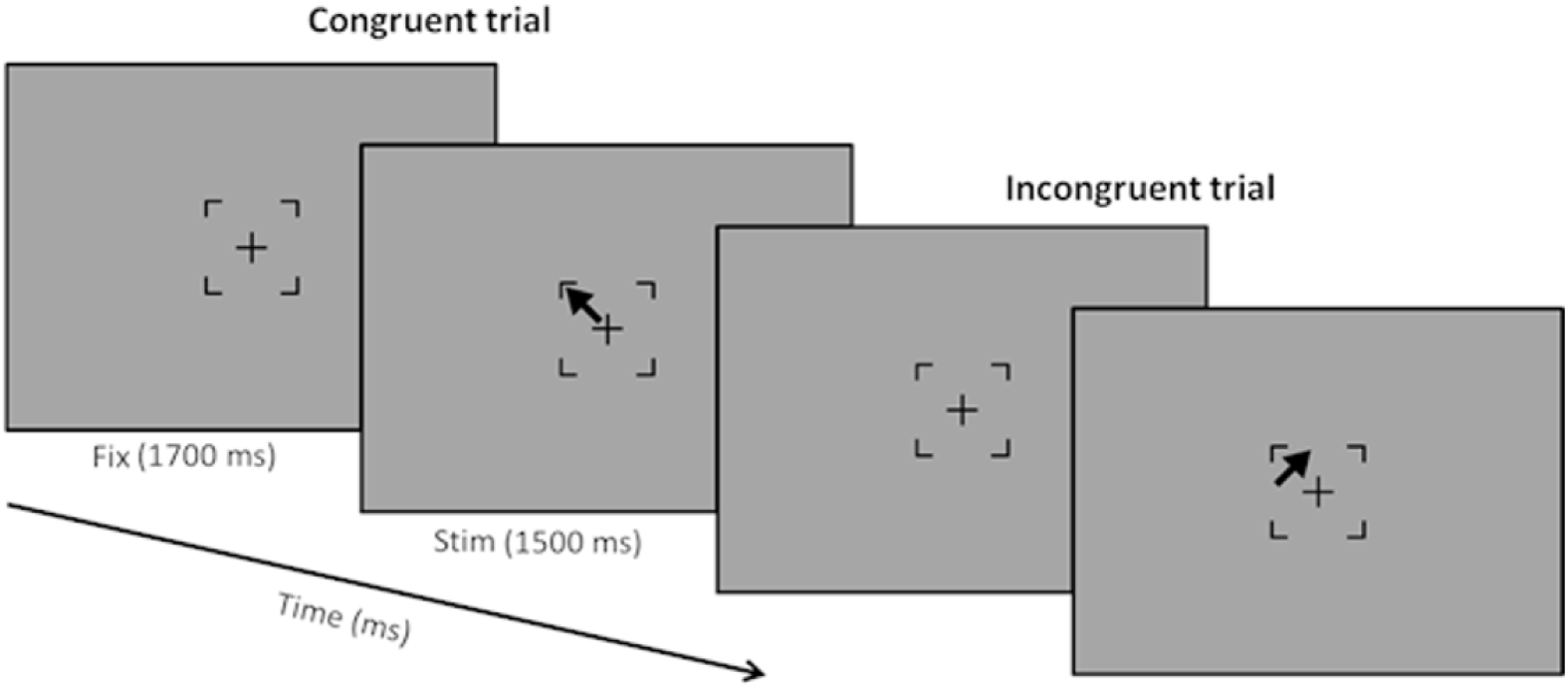
Perifoveal spatial Stroop task. Each trial began with a fixation cross (*Fix*, 1700 ms), followed by the stimulus (*Stim*, max 1500 ms). The stimulus consisted of an arrow pointing toward one of four quadrants (Target feature) and presented in one of the same four spatial locations (Distractor feature). A match or mismatch between arrow direction and position resulted in Congruent and Incongruent trials, respectively. Participants reported the arrow’s direction (Target) while ignoring its spatial position (Distractor). For a more extensive description, see Viviani et al. (2026)

Each trial began with a central fixation cross enclosed in the partial outline of black square (1700 ms), followed by the experimental stimulus which remained on screen for a maximum of 1500 ms. The experiment comprised 17 blocks of 40 trials each, plus an initial training block (16 trials) with performance feedback.

To induce proactive and predictive reactive stability, we manipulated the proportion congruency (PC) at two levels. Proactive stability was stressed via a list-wide PC (LWPC) manipulation, where the global probability of congruent trials varied across blocks (30%, 50%, or 70% of congruent trials). Predictive reactive stability was induced via an item-specific PC (ISPC) manipulation, where specific stimulus positions were associated with different conflict likelihoods within each block (PC could be 20%, 40%, 50%, 60% or 80%). Additionally, we manipulated Contingency (i.e., the conditional probability of a response given a stimulus position reflecting low-level stimulus-response associations), orthogonalizing it as much as possible from the PC manipulations to disentangle higher-order stability effects (i.e., induced by LWPC and ISPC) from low-level confounds (i.e., Contingency, see Schmidt, 2019). Although we manipulated these variables at the block-level, we have never used them in the analyses. As explained in detail below, we estimated them at the trial-level using an ideal Bayesian observer (to track the observer’s evolving belief about control demands) and we used these estimates in the analyses (see also Section S1.1, and Viviani et al., 2026 for a detailed description).

### 2.3. EEG data: Recording, Pre-processing, and single-trial ERSP data

The multivariate analyses reported here were performed on the same single-trial Event-Related Spectral Perturbation (ERSP) data previously computed for the univariate analyses reported in Viviani et al. (2026). No new pre-processing or time-frequency decomposition was performed for the present study: both the pre-processed EEG datasets and the resulting single-trial ERSP data were taken directly from that work and are openly available in our project repository of our previous work on the Open Science Framework (OSF) at https://osf.io/4df9s/. For the reader’s convenience, we summarise below the procedures that are relevant to the interpretation of the present multivariate analyses; full details regarding the exact pipeline are reported in Viviani et al. (2026).

EEG data had been recorded at 500 Hz from 64 Ag/AgCl electrodes (EASYCAP GmbH, Germany) using BrainAmp amplifiers (Brain Products, Munich, Germany). Offline pre-processing was conducted in MATLAB via ad-hoc EEGLAB scripts (version 14.1.2; Delorme & Makeig, 2004). Continuous data were filtered (zero-phase Hamming-windowed sinc FIR high-pass and low-pass filters, 0.1–45 Hz) and bad channels removed (via *clean_rawdata* function). Artifacts (e.g., eye movements, muscle activity) were identified using Independent Component Analysis (fastICA algorithm; Winkler et al., 2015) and removed through an automatic selection procedure based on IClabel classification, followed by visual inspection. Bad channels were interpolated using spherical splines (Perrin et al., 1989). Epochs containing residual artifacts were detected based on extreme values, improbability, and kurtosis criteria, and corrected via the *Trial by Trial* (TBT) plugin to reject or interpolate channels on a trial-by-trial basis (see Viviani et al., 2026, for detailed parameters and rejection rates). In addition to the resulting clean non-baseline-corrected stimulus-locked dataset, a response-locked dataset was created by locking the data to the response time for each trial.

Single-trial ERSPs had been computed from both non-baseline-corrected datasets using complex Morlet wavelet convolution, over a frequency range from 4 to 30 Hz (1-Hz resolution), with the number of cycles increasing linearly with frequency (from 3 to 12 cycles). To account for non-systematic fluctuations while preserving representational patterns, spectral power was normalized trial-by-trial relative to the average power of the entire epoch (from -1500 to 1500 ms for stimulus-locked and -500 to 300 ms for response-locked data). For the present study, the resulting ERSP data were downsampled to 50 Hz (one point every 20 ms) for computational constraints. These single-trial ERSP data served as the input to all multivariate searchlight analyses reported below. For a more detailed explanation of the ERSP computation, we refer the reader to Viviani et al. (2026).

### 2.4. Multivariate data analysis

#### 2.4.1. Rationale and general approach

All multivariate pattern analyses (MVPA) were implemented using an integrated, multi-feature searchlight approach. This method identifies representational spatio-temporo-spectral patterns within a combined multidimensional neural space defined by neighboring electrodes, time points and spectral frequencies, without collapsing informative dimensions.

We used this searchlight approach with Representational Similarity Analysis (RSA) to estimate, at each searchlight coordinate, the encoding strength of stability-related representations (i.e., task and conflict-expectancy), as well as whether —and in which direction— task representation encoding strength is modulated by trial-by-trial conflict-expectancy levels. Then, ridge-regression decoding, which allows predicting continuous variables rather than performing classification (Cohen et al., 2011), was applied to the same neural patterns to assess the decodability of the same informational content, thereby providing convergent validation of the RSA findings by testing whether the same information is structured in a format that could be read out by downstream processes, and isolating the most robust representational patterns. Specifically, a descriptive conjunction was performed between the resulting encoding and decoding maps to identify the specific spatio-temporo-spectral patterns where encoding (RSA) results were validated by successful functional decodability (Ridge). Specifically, we computed the intersection (overlap) between the cluster-corrected significant RSA statistical maps and the cluster-corrected significant ridge-regression decoding maps (RSA ∩ Ridge; see Sections 2.4.3-2.4.4).

To maximize sensitivity and representational precision, all MVPA analyses utilized a multilevel, trial-level approach. Rather than aggregating trials into categorical conditions, we modeled the trial-by-trial fluctuations of both task variables (Target and Distractor) and conflict-expectancy variables. To estimate the latter, we utilized the Hierarchical Gaussian Filter (HGF; Mathys et al., 2011) —an ideal Bayesian observer— to compute the trial-by-trial probability estimates of conflict at both the global (LWPC) and item-specific (ISPC) levels based on trial history (see below, for more details, see Di Pietro & Viviani, 2026, and for a tutorial, see https://osf.io/qmu7g/overview). Statistical inference was performed in two steps: first, multiple linear regressions were performed for each participant to estimate individual effects (representing encoding strength or decodability); second, group-level significance was assessed on the resulting subject-level coefficients. To handle the high dimensionality of the multivariate maps, all statistical results were corrected for multiple comparisons using a cluster-based permutation approach.

Using this framework, we tested our mechanistic representational hypotheses. In the post-stimulus period (both stimulus-locked and response-locked), we assessed: i) the encoding of task representations (i.e., Target and Distractor), expecting the Target representation to be significantly encoded to subserve effective performance; ii) the encoding of conflict-expectancy representations, expecting the information of LWPC and ISPC to be encoded sufficiently early —and with sufficient temporal precedence over the task representations they are hypothesized to modulate— to serve as functional inputs to downstream stability processes; iii) how global and item-specific conflict-expectancies modulate the encoding of the task representations, expecting that higher conflict expectations (i.e., lower LWPC/ISPC) enhance the Target encoding strength and reduce the Distractor encoding strength. To the best of our knowledge, for the first time we directly tested this directionally specific modulation, by modeling the interaction between task variables (Target/Distractor) and conflict-expectancy levels (LWPC/ISPC). In the pre-stimulus, where we specifically targeted the main effects of LWPC and ISPC representations, we determined whether conflict-expectancy information is encoded prior to the emergence of conflict.

The specific procedures for variable modeling, the implementation of the RSA and decoding analytical pipelines, and the information-theoretic metrics are described in the following sections, with further technical specifications and mathematical details provided in the Supplementary Materials (SM, available at https://osf.io/37b4e/overview; sections and figures therein are denoted by the prefix “S”, e.g., Section S1.3, Fig. S1).

#### 2.4.2. Experimental variables and representational models

Following the analytical approach described above, we modeled the informational content of both task and conflict-expectancy representations. Although the underlying theoretical models were consistent across analyses, the experimental variables were computed differently to meet the specific requirements of RSA and ridge-regression decoding. Full mathematical details on variable computation are provided in Supplementary Materials (Section S1.2) and all codes are available at https://osf.io/37b4e/overview.

##### Task variables: Target and Distractor

The task-relevant (Target) and task-irrelevant (Distractor) representations reflect the core dimensions on which stability should operate. In our paradigm, these correspond to the target arrow direction (DIR) and its spatial position (POS), respectively. Both features were defined by four possible levels (upper-left, upper-right, lower-right, and lower-left).

For RSA, these dimensions were modeled as categorical Representational Dissimilarity Matrices (RDMs) based on logical divergence. The Target RDM reflected trial-to-trial similarity when two trials shared the arrow target direction (i.e., the same required response), while the Distractor RDM reflected similarity when trials shared the same spatial position (i.e., the same irrelevant location). For further details see Section S1.2.1.1).

For ridge-regression decoding, we decomposed the Target and Distractor dimensions into their horizontal and vertical components, in order to have binary 2-D variables for each trial to decode. Crucially, to handle the intrinsic correlation between Target and Distractor dimensions (which overlap perfectly in congruent trials) and the other predictors, we applied a residualization procedure. This allowed us to isolate the unique informational content of each task dimension by regressing out the variance associated with the competing feature, trial congruency, and current conflict-expectancy estimates. This procedure ensured that the decoding performance for a specific dimension (e.g., Target) was not confounded by other task or demand-level factors (for further details, see Section S1.2.1.2).

##### Conflict-expectancy variables: LWPC and ISPC

Conflict-expectancy variables reflect the representations of trial-by-trial conflict probability manipulations (LWPC and ISPC) that drive continuous adjustments of proactive and predictive reactive stability. To account for the dynamic nature of stability, where expectations are continuously updated based on trial history, we used the Hierarchical Gaussian Filter (HGF; Mathys et al., 2011; see Viviani et al., 2026) as an ideal Bayesian observer to obtain trial-specific estimates of congruency (and thus conflict) probability.

LWPC is the global expectation of conflict that triggers proactive stability. We computed LWPC as the trial-by-trial probability of stimulus congruency estimated by an ideal observer (such that lower values reflect higher global conflict expectation; see Section S1.2.2.1). For RSA, we constructed the LWPC RDM by modeling the representational distance between trial-by-trial global conflict expectations (see Section S1.2.2.3). In addition, to test whether global conflict expectations modulate the encoding strength of the task representations in a directional manner, we built a dedicated trial-by-trial pairwise level matrix (PLM) encoding the shared LWPC level of each trial pair (LWPC_level; see Section S1.2.2.3), which was used to compute the interaction terms entered in the RSA model as modulators of the Target and Distractor RDMs. For example, in the LWPC_level PLM, a [0.1 / 0.1] pair is coded as 0.1 to reflect high stability demand, while a [0.1 / 0.9] pair is coded as 0.5 to cancel out the two opposite stability demands. This allows us to move beyond establishing that expectations are encoded, toward specifying how —and in which direction— they modulate task representations (see Section 2.4.3). For ridge-regression decoding, we used the trial-level estimates after a residualization procedure designed to isolate the unique LWPC variance (see Section S1.2.2.4).

ISPC is the expectation of conflict conditional to a specific item (POS) that triggers predictive reactive stability. We first estimated the full profile of trial-by-trial conflict probability for all four stimulus positions (ISPC_total). From this, we derived a trial-level ISPC estimate that isolated the representation of conflict probability for the specific stimulus position encountered in each trial relative to the others (i.e., ISPC_weighted; see Section S1.2.2.1). For RSA, we constructed RDMs for both ISPC_total (capturing the full latent landscape of item-specific conflict probabilities maintained prior to stimulus onset, used in the pre-stimulus analyses) and ISPC_weighted (capturing the weighted trial-level item-specific conflict expectations, the conflict expectation triggered once the actual stimulus location is identified on a given trial) (see Section S1.2.2.3). In addition, mirroring the approach used for LWPC, we built a dedicated PLM encoding the shared ISPC level of each trial pair (ISPC_level; see Section S1.2.2.3), which was used to compute the interaction terms entered in the RSA model as modulators of the Target and Distractor RDMs. For ridge-regression decoding, we used the residualized versions of ISPC_weighted and ISPC_total to assess their unique contribution to neural decodability (see Section S1.2.2.4).

#### 2.4.3. Representational Similarity Analysis (RSA) implementation

To characterize the strength of the neural encoding of stability representations, we investigated the similarity between the theoretically defined model matrices and the multivariate neural patterns (for full technical specifications of the RSA pipeline see Section S1.3 and Fig. 2).

**Figure 2.**
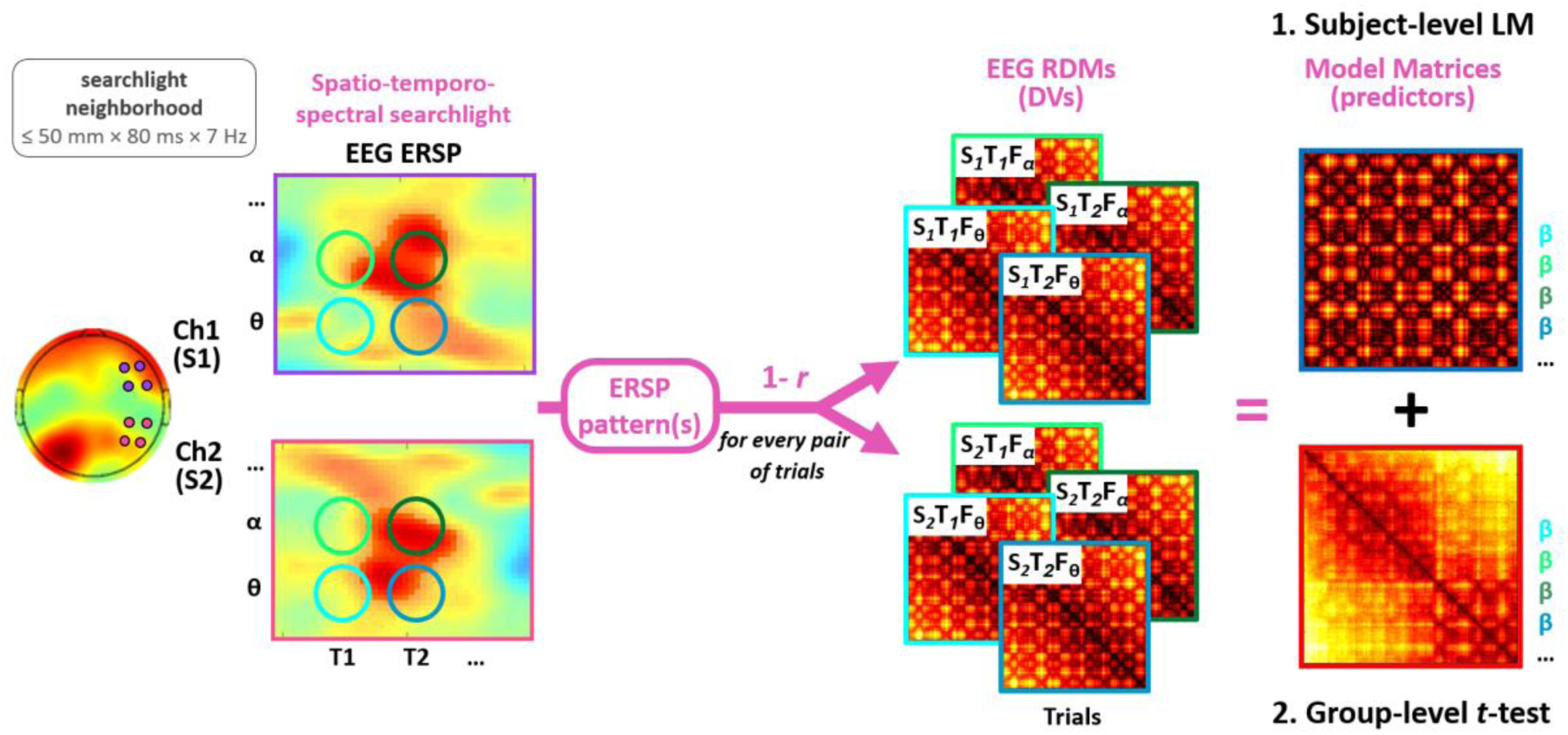
Multi-feature searchlight RSA pipeline. Illustration of the trial-level spatio-temporo-spectral searchlight Representational Similarity Analysis (RSA) framework. The pipeline operates on the participant’s event-related spectral perturbation (ERSP). A trial-level searchlight iterates across the space, time, and frequency dimensions of the ERSP: at every searchlight coordinate —defined by a spatial center (a channel S; e.g., S₁/S₂ centered on Ch₁/Ch₂), a temporal center (T; e.g., T₁/T₂), and a spectral center (F; e.g., Fα/Fθ)— a multidimensional neighborhood is drawn around that center, spanning a spatial radius of ≤ 50 mm, a temporal window of 80 ms, and a spectral window of 7 Hz (top-left callout). Within this neighborhood the ERSP is extracted trial by trial as an ERSP pattern. Then, the correlation distance (1 − Pearson’s r) between each of these ERSP patterns is computed for every pair of trials, yielding the brain RDM (dependent variable; “Trials” denotes both matrix axes). (1) At the subject level, multiple linear regressions predict brain RDMs using theoretically-defined model matrices as predictors —two are shown here as examples; the full model included dedicated RDMs for Target, Distractor, LWPC, ISPC, and low-level confounding, plus pairwise level matrix (PLM)-based interaction terms, which encode the shared conflict-expectation level of each trial pair and enter the regression as multiplicative modulators of the Target and Distractor RDMs (see Section 2.4 and Section S1.3 in Supplementary Materials). This yields one beta (β) map per spatio-temporo-spectral pattern (color-coded to match brain RDMs) which index the encoding strength of each predictor at each spatio-temporo-spectral pattern (where, when, and in which band that predictor contributes to explaining the brain RDMs). (2) At the group level, one-sample t-tests are performed on the β maps across participants, with directional contrasts defined a priori to test the hypothesized representational reconfigurations.

##### Brain RDM computation

For each participant, we computed trial-by-trial spatio-temporo-spectral brain RDMs using the searchlight approach. Specifically, at each time point, frequency, and channel, the brain RDM was calculated as the correlation distance (1 − Pearson’s r) between the ERSP patterns across every pair of trials, within a multidimensional neighbor-space. This space was defined by a spatial radius (Euclidean distance <= 50 mm), a temporal window of 80 ms (corresponding to 5 time points), and a spectral window of 7 Hz (see Fig. 2). Brain RDMs were computed for: (i) stimulus-locked pre-stimulus data (–1000 to 0 ms); (ii) stimulus-locked post-stimulus data (0 to 1000 ms); and (iii) response-locked data (–500 to 300 ms relative to response time).

##### RSA regression and group-level inference

As described in Section 2.4.1, we used a condition-rich RSA approach, in which both the brain RDMs and the model matrices were computed trial-by-trial and then entered into a trial-level two-step multiple regression approach at the whole-brain level. At the first level, we performed multiple linear regressions for each participant to predict the observed brain RDMs at each searchlight coordinate (i.e., multivariate pattern) using the model matrices as predictors (see Section S1.3.1). This allowed us to estimate the representational strength (beta values) with which each model was encoded.

For the post-stimulus and response-locked analyses, we tested the following regression model:

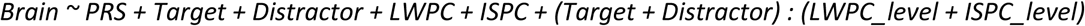

This model served two purposes. First, it assessed how the task and conflict-expectancy representations were encoded, through the main effects of the corresponding RDMs (i.e., Target, Distractor, LWPC, and ISPC; note that here ISPC refers to the weighted one, namely the specific expectancy triggered by the stimulus). Second, and central to our hypotheses, it tested how conflict-expectancy representations modulated the encoding strength of the task representations, through the interaction terms, which were built using the PLMs of the conflict expectations (LWPC_level and ISPC_level; see Sections S1.2.2.3 and S1.3). By encoding where each pair of trials lies on the expectancy scale, the PLMs make it possible to establish whether higher conflict expectation strengthens the encoding of the Target and/or weakens the encoding of the Distractor. Moreover this model assessed whether conflict expectations reconfigured the neural code of task dimensions while also controlling for low-level contingency (PRS; Schmidt, 2019; Viviani et al., 2026, see Section S1.2.2.2).

To ensure that the multiple regression RSA model yielded stable and unconfounded parameter estimates, we formally assessed multicollinearity across all vectorized predictor matrices (main effect RDMs and interaction terms). Pairwise correlations between model predictors were minimal (all |*r*| ≤ .20, mean |*r*| = .046), and collinearity diagnostics revealed negligible variance inflation across the entire predictor set, with Variance Inflation Factors (VIFs) remaining near optimal baseline (mean VIF = 1.06, maximum VIF < 1.08). These diagnostics confirm that the estimated partial regression weights (beta coefficients) reliably reflect the unique, non-overlapping representational variance of each computational model and interaction without distortion from collinearity.

For the pre-stimulus analysis, where task dimensions are unknown, the model focused on the anticipatory representation of global and item-specific conflict expectations:

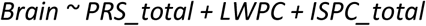

Here, we utilized the total probability distributions (i.e., ISPC_total and PRS_total) based on multinomial distributions to investigate whether the brain encodes the full configuration of item-specific conflict expectations before knowing which stimulus feature will appear in the upcoming trial. Multicollinearity was not problematic in this case either (all |*r*| ≤ .51, mean |*r*| = .25; mean VIF = 1.30, maximum VIF < 1.45).

At the second level, group-level significance was assessed via one-sample t-tests on the resulting participant-specific beta maps. One-tailed tests were used throughout, as all effects had a directional a priori hypothesis (Montefinese et al., 2025). For the main effects (i.e., RDMs), encoding strength was expected to be positive, since representational similarity matrices are bounded to quantify positive pattern matching; negative beta coefficients in model RDMs lack theoretical and neurocomputational interpretability. For the interaction terms (i.e., PLM-dependent modulation of RDM encoding strength), directional contrasts were formally defined based on our explicit a priori hypotheses regarding representational reconfiguration (i.e., testing specifically for Target enhancement and Distractor suppression under high conflict expectations): since lower PLM values index higher conflict expectations, these correspond to negative and positive interaction betas, respectively, and the sign of the former was inverted before one-tailed testing.

The resulting statistical parametric maps (see Section S1.3.2) were corrected for multiple comparisons using a cluster-based permutation approach based on the *tmass* statistics, using 2000 permutations to estimate the null distribution.

#### 2.4.4 Ridge-regression decoding implementation

To complement the RSA and assess the decodability of task and conflict-expectancy informational content, we performed ridge-regression decoding (see Section S1.4 for technical details).

##### Decoding procedure and searchlight parameters

For each participant, we implemented a spatio-temporo-spectral searchlight decoding approach. Similar to the RSA, the analysis targeted single-trial ERSP values within a multidimensional neighbor-space, which served as the design matrix for predicting the experimental variables (Euclidean distance < 50 mm; 80-ms temporal window). Based on the spectral dynamics observed in previous univariate analyses (Viviani et al., 2026), we targeted five frequency bands: Theta (4–7 Hz), Alpha (8– 12 Hz), and three Beta sub-bands (Beta1: 13–18 Hz; Beta2: 19–24 Hz; Beta3: 25–30 Hz). At each searchlight coordinate, a ridge-regression model was trained and tested using a 5-fold balanced cross-validation procedure. The decoding performance was indexed by the cross-validated *R^2^*, reflecting the proportion of variance in the (unique) experimental variables explained by the multivariate neural patterns.

##### Decoding models

To ensure that the decodability reflected unique informational content, we used the residualized versions of the task features (Target, Distractor) and conflict-expectancy estimates (LWPC, ISPC) as the predicted variables (see Section 2.4.2, and see Sections S1.2.1.2 and S1.2.2.4, for the residualization pipeline). The models for the post-stimulus and response-locked analyses for the main effects allowed us to identify which spatio-temporo-spectral EEG features can decode the task and conflict-expectancy predictors. Moreover, by testing also the interaction terms (e.g., Target x LWPC), we also assessed the decodability of the interplay between conflict-expectancy and task variables. Pre-stimulus analyses targeted the decodability of global and item-specific conflict expectations (LWPC and ISPC_Total).

##### Statistical inference

Group-level significance was assessed by performing a one-sample non-parametric test on the subjects’ *R^2^* values at each searchlight coordinate, comparing them against a null distribution generated through a permutation procedure (1000 permutations; see Section S1.4.2). This approach allowed us to determine the probability of obtaining the observed decoding performance by chance, while preserving the spatio-temporo-spectral structure of the EEG signal. The resulting statistical parametric maps were corrected for multiple comparisons using a cluster-based permutation test (2000 permutations; see Section S1.4.3 for technical details).

## 3. Results

In the following sections, we report the results of the RSA analyses. Given that ridge-regression decoding was employed as a functional validation tool rather than as a primary analysis, decoding results are not reported independently. Instead, we highlight the subset of RSA findings that were convergently confirmed by significant decoding performance —identified via descriptive conjunction (i.e., statistical map intersection)— and thus treated as the strongest and most valid representational patterns. Accordingly, the plots display the spatio-temporo-spectral RSA results that survived cluster-based correction, with light green outlines superimposed to indicate the conjunction with the decoding results (see Table 1 for a summary of all results).

**Table 1.** – Summary of findings.

| Domain | Effect | Section | Fig. | Epoch | Spatial Pattern | Time Wind. (Peak) in ms | Frequency Band | Mechanistic and Computational Role |
|---|---|---|---|---|---|---|---|---|
| **Task** | Target | 4.1.1 | S1 | STIM | R. F+C | 20 - 580<br>(420) | Upper-Beta | **Action-oriented goal representation**:<br>Direct encoding of task rule and response choice (arrow direction → motor output), driving S–R translation for action selection. |
| **Task** | Distractor | 4.1.2 | S2 | STIM | R. F+C | 0 - 1000<br>(200) | Lower-Beta | **Automatic capture & motor migration**:<br>Early automatic encoding of task-irrelevant sensory representation (position), migrating into motor circuits generating conflict at response execution. |
|  |  |  | S8 | RESP | M. C | –480 - –260<br>(–400) | Upper-Beta |  |
| **Expect.** | LWPC | 4.2.1 | 3 | STIM | R. F+C<br>L. P+O | 0 - 420<br>(240) | Theta | **Early and sustained encoding of global conflict-expectancy representation**:<br>Dynamic Bayesian (history-dependent) representation, generated by tracking of ongoing conflict evidence and integrated into memory, continuously maintained across the trial cycle. Response-locked recruitment of Alpha-mediated gating supports actionable control at execution. |
|  |  |  | S9 | RESP | L. Wide | –500 - 300<br>(–220) | Theta + Alpha |  |
| **Expect.** | LWPC | 4.2.2 | S3 | STIM | M. FC<br>L. P<br>R. T | 660 - 1000<br>(800) | Lower-Beta | **Anticipatory trial-by-trial cycle of global conflict-expectancy representation**:<br>1) Post-stimulus updating to incorporate trial outcome into the belief state;<br>2) ITI maintenance in a protected state;<br>3) Pre-stim reinstatement into an actionable top-down predictive state. |
|  |  |  | 7 | PRE | R. FT | –1000 - –480<br>(–680) | Upper-Beta |  |
|  |  |  | 7 | PRE | M. FC | –540 - –140<br>(–420) | Lower-Beta |  |
| **Expect.** | ISPCw | 4.2.3 | S4 | STIM | L. PO | 40 - 380<br>(160) | Alpha | **Early stimulus-triggered encoding of item-specific conflict-expectancy representation**:<br>Sensory-anchored, dynamic Bayesian (history-dependent) representation, initially expressed as an early attentional gating on visuo-spatial processing, then reinstated as an endogenous prior representation to guide conflict anticipation prior to response execution. |
|  |  |  | 5 | RESP | M. PO | –380 - 80<br>(–260) | Lower-Beta |  |
| **Expect.** | ISPCt | 4.2.4 | 8 | PRE | L. PF | –1000 - –660<br>(–920) | Upper-Beta | **Anticipatory trial-by-trial cycle of item-specific conflict-expectancy representation**:<br>1) ITI maintenance of the full item-specific conflict probability landscape in a protected state;<br>2) Pre-stim reinstatement of the maintained landscape, ready for stimulus-triggered retrieval. |
|  |  |  | 8 | PRE | F+C<br>P+PO | –300 - 0<br>(–220) | Theta + Alpha |  |
| **Proactive** | Target<br>x LWPC | 4.3.1 | 4 | STIM | L. C+T+P | 0 - 60<br>(20) | Lower-Beta | **Early and Sustained proactive strengthening of the Target representation:**<br>High global conflict expectations increase Target strength via early predictive routing boosting task-relevant sensory processing, sustained gating shielding the Target from competing inputs, and a pre-response reinstatement to guide execution. |
|  |  |  | 4 | STIM | L. F | 80 - 180<br>(120) | Alpha |  |
|  |  |  | 4 | STIM | M. P+PO | 360 - 620<br>(480) | Alpha |  |
|  |  |  | S10 | RESP | M. PO | –160 - –100<br>(–140) | Lower-Beta |  |
| **Proactive** | Distractor<br>x LWPC | 4.3.2 | S5 | STIM | R. PF | 500 - 540<br>(520) | Upper-Beta | **Late and circumscribed proactive weakening of the Distractor representation**:<br>High global conflict expectations reduce Distractor strength via a late prefrontal executive brake actively suppressing the interfering spatial representation, and a pre-response weakening of its memory trace. |
|  |  |  | S11 | RESP | L. T | –360 - –240<br>(–320) | Theta + Alpha |  |
| **Predictive Reactive** | Target<br>x ISPC | 4.4.1 | S6 | STIM | R. CP+P | 400 - 520<br>(460) | Theta | **Late and response-anchored item-triggered predictive reactive strengthening of the Target representation**:<br>High item-specific conflict expectations increase Target strength via attentional prioritization of the task-relevant code, and a reinstatement to impose a pre-response bias. |
|  |  |  | S12 | RESP | R. FT | –220 - –100<br>(–160) | Lower-Beta |  |
|  |  |  | S12 | RESP | R. P | –80 to 0<br>(–60) | Lower-Beta |  |
| **Predictive Reactive** | Distractor<br>x ISPC | 4.4.2 | S7 | STIM | L. P+PO | 240 - 400<br>(300) | Lower-Beta | **Early item-triggered predictive reactive weakening of the Distractor representation**:<br>High item-specific conflict expectations reduce Distractor strength via perceptual dampening through direct spectral gating and attentional gating, and a pre-response direct motor braking. |
|  |  |  | 6 | RESP | R. PO | –480 - –140<br>(–320) | Theta + Alpha |  |
|  |  |  | 6 | RESP | L. P | –120 - –40<br>(–100) | Upper-Beta |  |

**Note**. Time windows refer to the full extent of each significant RSA cluster (peak latency in brackets); spatial patterns and frequency bands refer to the strongest encoding pattern (i.e., the RSA ∩ Ridge conjunction). See Section 3 for the full spatial and spectral extent of each cluster.

For each result, we first describe the RSA-defined spatio-temporo-spectral cluster, and then specify what we will call the “strongest” encoding pattern within it (or “strongest modulation effect” for interaction terms) —i.e., the convergent encoding pattern where RSA was also validated by successful ridge-regression decoding. When the strongest encoding pattern is spatially, temporally, or spectrally narrower than the broader significant RSA cluster, we report its specific spatio-temporo-spectral coordinates; when it entirely overlaps with the RSA cluster, we do not repeat the shared features.

Results are reported in three sections: stimulus-locked (Section 3.1), response-locked (Section 3.2), and pre-stimulus (Section 3.3). Given our focus on anticipatory and stimulus-driven mechanisms, we restrict the reported findings to patterns emerging within a latency window relevant for the aims of the study: to specifically isolate representational dynamics driving decision-making and action selection, we excluded post-evaluative effects occurring “exclusively” later than 700 ms (stimulus-locked) or emerging exclusively after motor execution (response-locked), unless directly relevant to the mechanistic interpretation of the findings (e.g., the late LWPC main effect cluster in Section 3.1).

Across all analyses, main effects indicate where, when, and in which frequency band a given model matrix was significantly encoded, reflecting the strength with which each representation was expressed in the neural patterns. In the stimulus- and response-locked analyses, we additionally tested interaction terms capturing the modulation of task representations (Target, Distractor) by conflict-expectancy levels (LWPC_level, ISPC_level). As detailed in Section 2.4.3, these interactions were tested with directional a-priori hypotheses following our theoretical framework: higher conflict expectations (i.e., lower LWPC/ISPC values) were expected to enhance the encoding strength of the Target representation and reduce the encoding strength of the Distractor representation. Because both directions were tested one-tailed against the hypothesized effect, a significant interaction cluster appears with the same positive sign in the statistical maps (i.e., red in the plots) for both the Target and the Distractor interactions, despite reflecting empirically opposite modulations of encoding strength. In other words, red clusters in the Target × LWPC/ISPC maps indicate that higher conflict expectations strengthen the Target representation, whereas red clusters in the Distractor × LWPC/ISPC maps indicate that higher conflict expectations weaken the Distractor representation.

All tested models also included predictors for Contingency confounder, but its effects will not be reported here, as they lie beyond the scope of the present study.

Despite the stringent statistical corrections required by the high-dimensional spatio-temporo-spectral search space, the analyses still yielded a very large number of valid significant clusters. Specifically, all reported representational geometries successfully survived our rigorous two-tier validation framework, which ensures that they reflect robust, cross-methodologically validated neural information capable of out-of-sample generalization: (1) cluster-corrected significant RSA results (*p*(encoding) < .05; whole-brain cluster-based permutation test, t*mass* statistic evaluated against 2000 permutations; see Section S1.3.2) and (2) statistical map intersection (conjunction) with cluster-corrected significant results of the independent, cross-validated ridge-regression decoding (*p*(decoding) < .05; see Section S1.4.3). To provide a structured and readable narrative of these high-dimensional results without omitting data, we complemented the predefined latency window defined above with a standard extent reporting threshold: we focus in the main text on primary peak clusters whose footprint exceeds 25 contiguous spatio-temporal points —a criterion ensuring that reported patterns span beyond the minimal volume of a single searchlight kernel (80 ms × 50 mm) and represent coherent neural dynamics rather than localized edge transients. For each theoretically defined model and interaction, we report in the main text the primary peak clusters and their dominant spatio-temporo-spectral trajectory. To limit the length of the main text, selected figures are reported in the Supplementary Materials, available at https://osf.io/37b4e/overview, referred to throughout the text with the prefix “S” (e.g., Fig. S1).

### 3.1. Stimulus-locked representational results

RSA on stimulus-locked spatio-temporo-spectral patterns showed that all theory-based representational models of interest we tested were significantly encoded. Moreover, all these predictors showed significant decodability through multivariate neural patterns with a high degree of overlap with those revealed by RSA encoding them.

#### 3.1.1. Task representations

The Target representation was significantly encoded by spatio-temporo-spectral neural patterns immediately after stimulus onset (20 ms) till 580 ms, over fronto-centro-posterior electrodes, right lateralized, and involving Beta1 and Beta3 frequencies. The strongest encoding pattern was observed at around 420 ms over right fronto-central electrodes and involving Beta 3 frequencies (Fig. S1). The Distractor was significantly encoded by right (pre)frontal and central multivariate patterns, sustained from stimulus onset up to 1000 ms, through Beta and later Alpha spectral patterns. The strongest encoding pattern was at around 200 ms, involving mainly Beta1-2 frequencies, and was significant again at later time windows (Fig. S2).

#### 3.1.2. Conflict-expectancy representations

The representation of the global conflict expectation (LWPC) was significantly encoded already at stimulus onset until 420 ms by left posterior and right (pre-)fronto-central spatial patterns, involving Theta and Alpha frequencies. The strongest encoding pattern peaked at around 240 ms through Theta spectral patterns (Fig. 3). LWPC representation was also encoded later in time (660-1000 ms) by mid fronto-centro-posterior and right temporal spatial patterns, involving Beta1 frequency. Here, the strongest encoding emerged at around 800 ms (Fig. S3).

**Figure 3.**
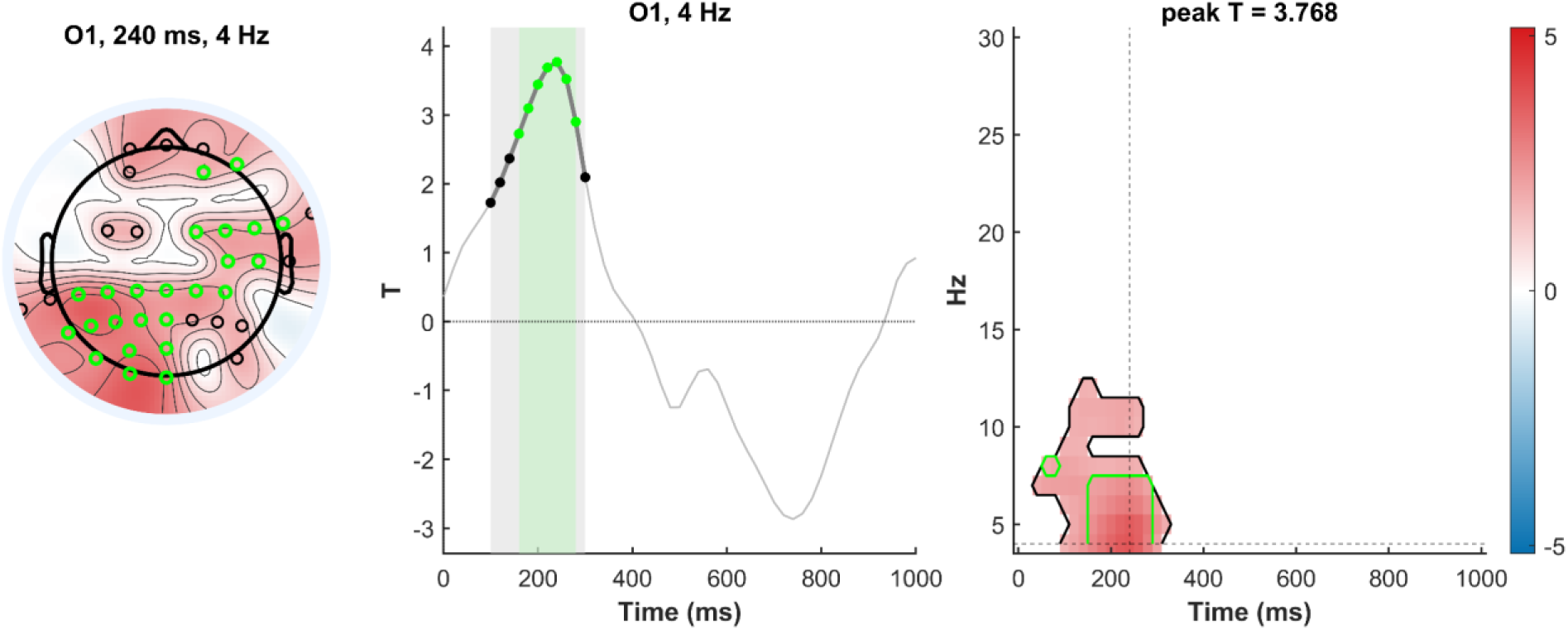
Early stimulus-locked representation of global conflict expectancy (LWPC). **(Left)** Topoplot showing the scalp topography of the representational encoding strength (*t*-statistic) at the peak spatio-temporo-spectral coordinate (indicated in the subplot title) of the significant RSA cluster (cluster-corrected *p* < .05, cluster-based permutation test). Black circles indicate electrodes contributing to the significant multidimensional cluster, while the subset of green circles indicate the electrodes contributing to the statistical map intersection with the cluster-corrected significant Ridge-regression decoding (RSA ∩ Ridge; descriptive conjunction). **(Middle)** Time course of the *t*-statistic extracted at the peak electrode and peak frequency (indicated in the subplot title) in the analyzed epoch. The gray shaded window and black timepoints indicate the temporal extent of the significant RSA cluster, while the subset of green shading and timepoints indicate the temporal extent of the RSA ∩ Ridge conjunction. **(Right)** Time-frequency map of the *t*-statistic at the peak electrode. The black contour indicates the boundary of the significant spatio-temporo-spectral RSA cluster, while the green contour highlights the sub-cluster from the RSA ∩ Ridge conjunction. Dashed vertical and horizontal lines mark the latency and frequency at the peak statistic (indicated in the subplot title). The colorbar on the right indicates *t*-values, sharing the same scale across both the scalp topography and the time-frequency map.

The item-specific conflict expectation representation (ISPC)— here referring to the ISPC of the stimulus position actually observed on each trial, weighted relative to the ISPC values of the other positions (see Section 2.4.2)— was encoded early as well, but slightly later than stimulus onset (40-380 ms), by Alpha bilateral posterior multivariate patterns. The strongest encoding pattern emerged at around 160 ms in left posterior spatial patterns (Fig. S4).

#### 3.1.3. Proactive modulation of Task representations

Global conflict expectation (LWPC) modulated the encoding strength of the Target: when it was higher (low LWPC), the Target representation became stronger. This interaction was expressed across three sub-clusters, all left-lateralized. An early sub-cluster was present already at stimulus onset (0–60 ms, peaking at 20 ms) over left centro-posterior electrodes, involving Beta1-2 frequencies. A second sub-cluster emerged shortly after, between 80 and 180 ms (peaking at 120 ms), over left fronto-central electrodes and involving Alpha frequencies. A third and later sub-cluster was expressed between 360 and 620 ms over medial parieto-occipital electrodes, again through Alpha frequencies, where the strongest modulation effect of the whole interaction was observed at around 480 ms (Fig. 4).

**Figure 4.**
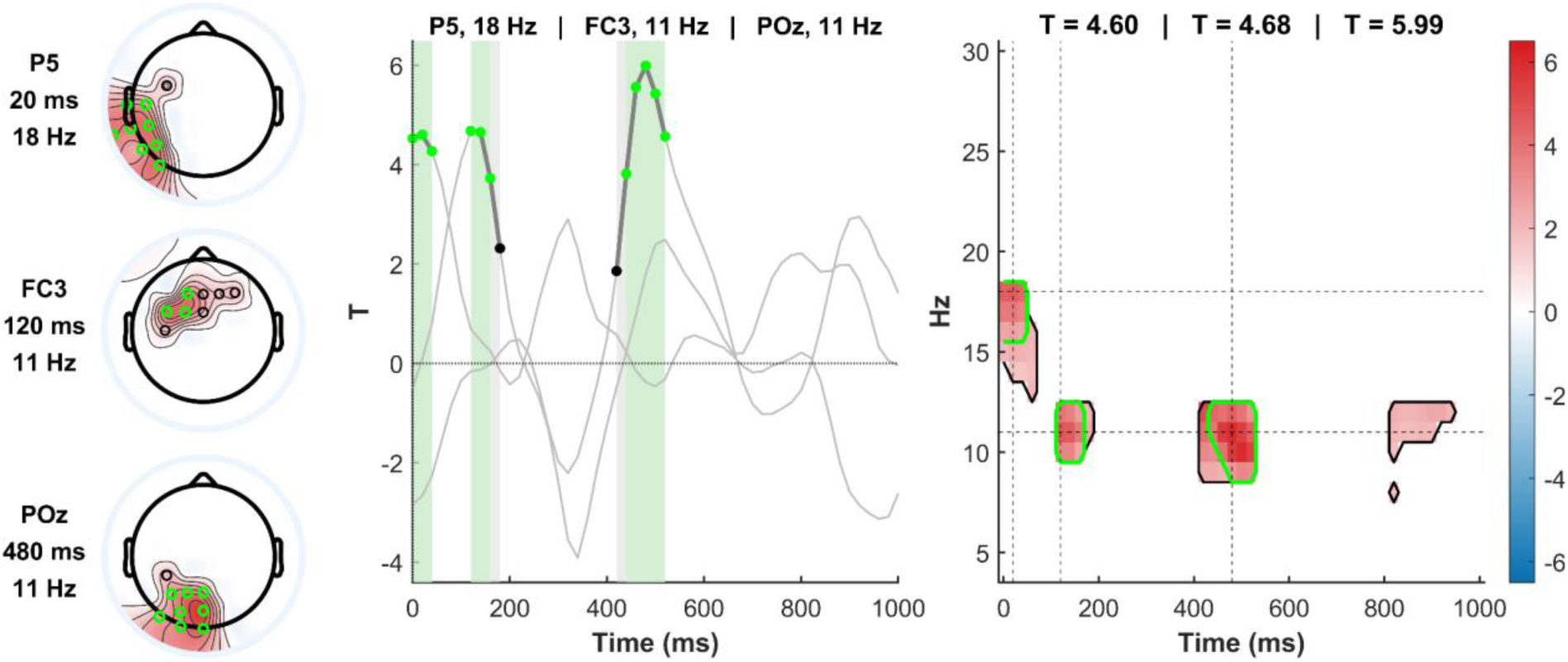
Stimulus-locked proactive modulation of Target representation by global conflict expectancy (Target x LWPC). Same format and conventions as in Figure 3, illustrating here the three distinct sub-clusters comprising the significant interaction. **(Left)** Scalp topographies at the three peak spatio-temporo-spectral coordinates. **(Middle)** Overlaid time courses of the t-statistic for each of the three peak channel-frequency pairs across the analyzed epoch. Shaded green areas and green dots indicate the respective temporal windows surviving the RSA & Ridge conjunction for each sub-cluster. **(Right)** Time-frequency map at the peak electrode, showing the three sub-clusters, with peak coordinates indicated by dashed lines and their respective peak t-values labeled above.

Global conflict expectation also modulated the encoding strength of the Distractor, though in a more circumscribed pattern: when the global conflict expectation was higher (low LWPC), the Distractor representation became weaker. This interaction was expressed between 500 and 540 ms over right pre-frontal spatial patterns, involving Beta3 frequencies, and peaked at around 520 ms (Fig. S5).

#### 3.1.4. Predictive reactive modulation of Task representations

Item-specific conflict expectation (ISPC) modulated the encoding strength of the Target representation: when it was higher (low ISPC), the Target representation became stronger. This interaction was expressed between 400 and 520 ms over right centro-parietal spatial patterns, involving Theta frequencies. The strongest modulation effect emerged at around 460 ms (Fig. S6).

ISPC also modulated the encoding strength of the Distractor: when it was higher (low ISPC), the Distractor representation became weaker. This interaction was expressed from 240 to 400 ms by Beta1 left parietal patterns. The strongest modulation effect emerged at around 300 ms (Fig. S7).

### 3.2. Response-locked representational results

RSA on response-locked spatio-temporo-spectral patterns showed that all theory-based representational models of interest, except for Target, were significantly encoded. Moreover, all these predictors could be decoded through multivariate neural patterns with a high degree of overlap with those revealed by RSA encoding them.

#### 3.2.1. Task representations

The Distractor was significantly encoded before response (from -480 to -260 ms) by mid-central spatial patterns, involving Beta2-3 spectral patterns. The strongest encoding pattern emerged at around -400 ms (Fig. S8).

#### 3.2.2. Conflict-expectancy representations

LWPC representation was sustainedly encoded from -500 to 300 ms around response by bilateral fronto-centro-posterior spatial patterns, involving Theta and Alpha frequencies, with the strongest encoding pattern emerging at around 220 ms before response (Fig. S9). ISPC representation was encoded from - 380 to 80 ms by medial posterior spatial patterns, involving Beta1-2 frequencies. The strongest encoding pattern emerged at around -260 ms, involving Beta1 frequencies (Fig. 5).

**Figure 5.**
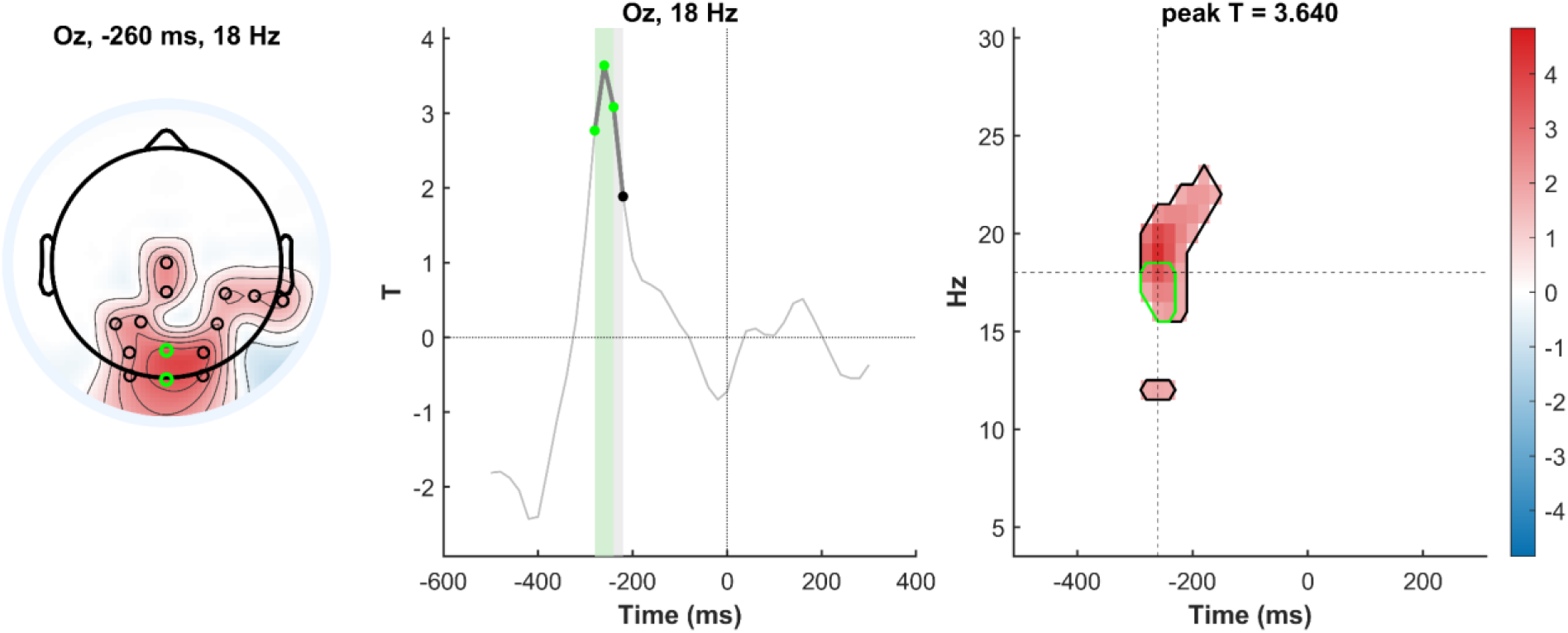
Response-locked representation of item-specific conflict expectancy (ISPC). Same format and conventions as in Figure 3, here with neural dynamics locked to response execution (time 0 ms).

#### 3.2.3. Proactive modulation of Task representations

Global conflict expectation (LWPC) modulated the encoding strength of the Target already before response: when the global conflict expectation was higher (low LWPC), the Target representation became stronger. This interaction was expressed between -160 and -100 ms over medial posterior electrodes, where the strongest modulation effect was observed at around -140 ms (Fig. S10). LWPC also modulated the encoding strength of the Distractor before response: when the global conflict expectancy was higher (low LWPC), the Distractor representation became weaker. This interaction was expressed from -360 to -240 ms over left temporal spatial patterns, involving Theta and Alpha frequencies. The strongest modulation effect emerged at around -320 ms (Fig. S11).

#### 3.2.4. Predictive reactive modulation of Task representations

Item-specific conflict expectation (ISPC) modulated the encoding strength of the Target already before response: when it was higher (low ISPC), the Target representation became stronger. This interaction was expressed across two temporally distinct sub-clusters, both right lateralized. The modulation emerged first between -220 and -100 ms before response over right fronto-temporal electrodes and involving Beta2 frequencies, then re-emerged between -80 and 0 ms over right centro-parietal electrodes and involving Beta1 frequencies. The strongest encoding modulation effects were observed at around -160 ms, and -60 ms (Fig. S12). Item-specific conflict expectation (ISPC) also modulated the encoding strength of the Distractor before response: when it was higher (low ISPC), the Distractor representation became weaker. This interaction was expressed across two temporally distinct sub-clusters. The modulation emerged first between -480 and -140 ms before response over right posterior electrodes and involving Theta and Alpha frequencies, then re-emerged between -120 and -40 ms over left parietal electrodes and involving Beta3 frequencies. The strongest modulation effects were observed at around -320 ms and -100 ms (Fig. 6).

**Figure 6.**
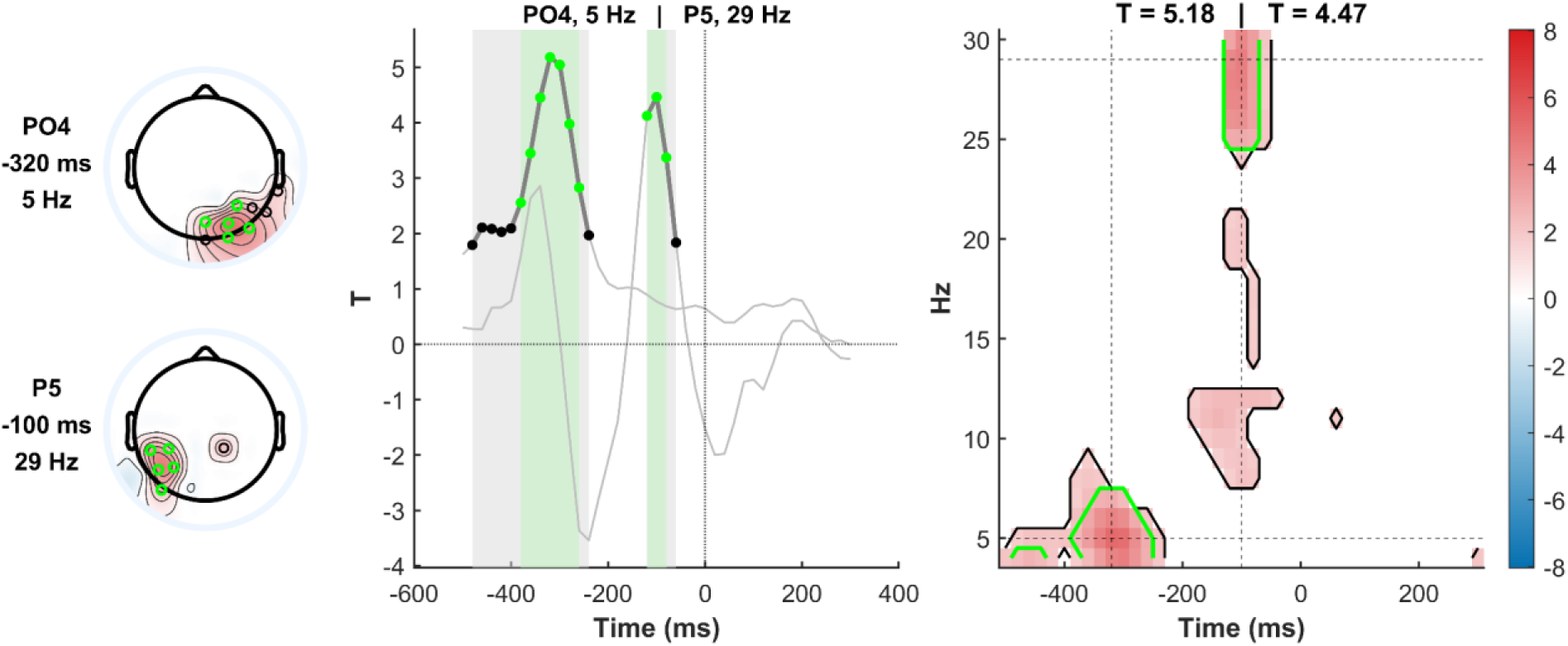
Response-locked predictive reactive modulation of Distractor representation by item-specific conflict expectancy (Distractor x ISPC). Same format and conventions as in Figures 3 and 4, here with data aligned to response execution (time 0 ms) and illustrating the two distinct sub-clusters comprising the significant interactions.

### 3.3. Pre-stimulus representational results

RSA on pre-stimulus spatio-temporo-spectral pattern revealed that both conflict-expectancy representations were significantly encoded, and that their decodability was significant and consistently overlapped with the multivariate neural patterns identified by RSA.

#### 3.3.1. Conflict-expectancy representations

LWPC representation was encoded across two pre-stimulus clusters. The first cluster emerged early, from -1000 to -480 ms before stimulus onset, over right anterotemporal spatial patterns and involving Beta frequencies, with the strongest encoding pattern observed at around -700 ms and peaking at 20 Hz. The second cluster emerged later, from -540 to -140 ms, over medial fronto-central spatial patterns and involving Beta1-2 frequencies, with the strongest encoding pattern at around -420 ms in Beta1 frequencies (Fig 7).

**Figure 7.**
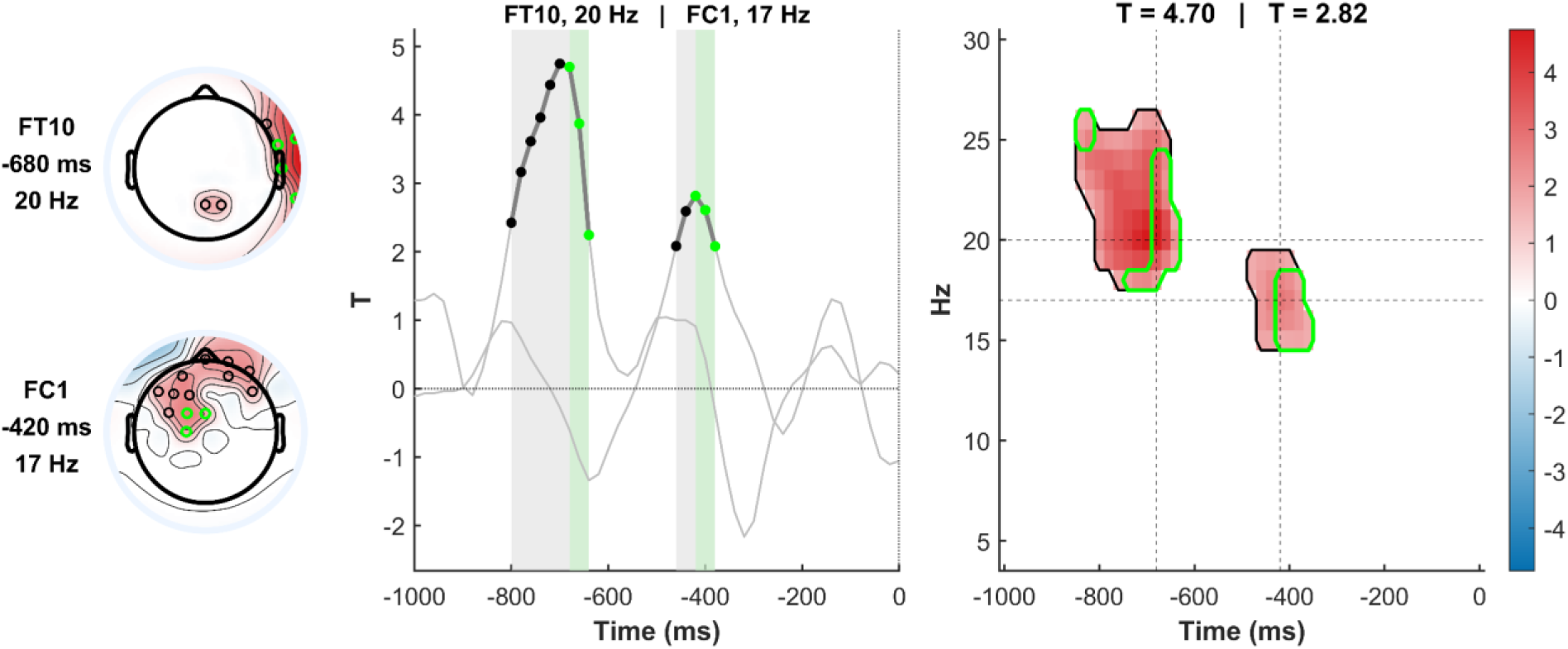
Pre-stimulus representation of global conflict expectancy (LWPC). Same format and conventions as in Figures 3 and 4, here with data locked to the pre-stimulus window (time 0 ms denotes stimulus onset) and illustrating the two distinct sub-clusters comprising the significant encoding.

The representation of the item-specific conflict expectation —here referring to the full configuration of item-specific conflict probabilities across all four possible stimulus positions (ISPC_total; see Section 2.4.2)— was encoded across two pre-stimulus clusters. The first cluster emerged early, from -1000 to -660 ms before stimulus onset, over left anterofrontal spatial patterns and involving Beta2-3 frequencies. The strongest encoding pattern was observed at around -920 ms over left prefrontal electrodes. The second cluster emerged later, from -300 to 0 ms, over widespread bilateral fronto-centro-posterior spatial patterns and involving Theta and Alpha frequencies, with the strongest encoding pattern at around -220 ms (Fig 8).

**Figure 8.**
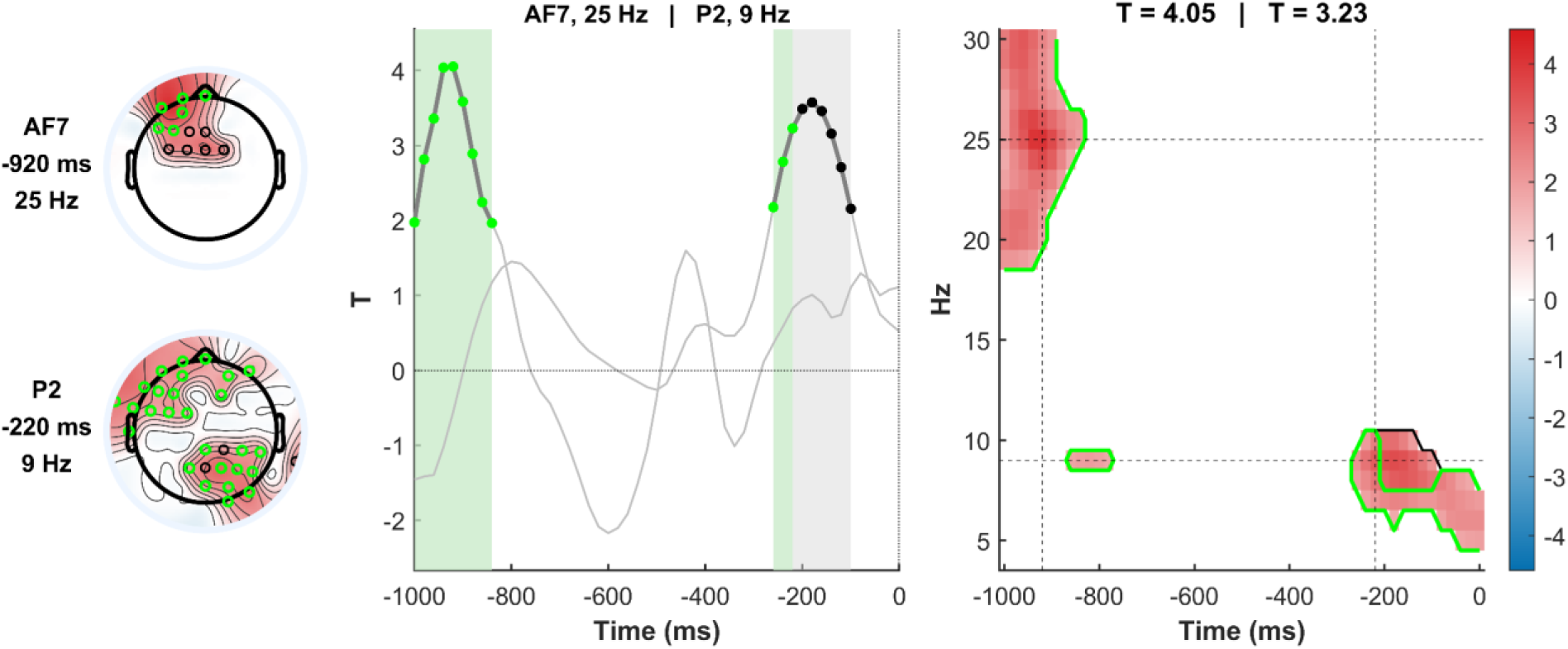
Pre-stimulus representation of the full item-specific conflict expectancy landscape (ISPC_total). Same format and conventions as in Figure 7.

## 4. Discussion

Cognitive stability is a core capacity of human cognition, enabling task focus through the prioritization of currently relevant tasks over competing ones (Egner, 2023). Beyond corrective reactive mechanisms that resolve conflict once it has emerged, stability also operates predictively, dynamically adjusting its level to match contextual demands through the statistical learning of conflict regularities, thereby anticipating conflict and minimizing its impact (Egner, 2023; Jiang et al., 2014). This predictive regulation can be implemented either proactively —sustainedly upregulating task focus when high global conflict is expected (as postulated by the Dual Mechanisms of Control; Braver, 2012)— or, as we recently proposed, through a predictive reactive mode that leverages item-specific conflict expectations to prepare the system before conflict fully emerges (Viviani et al., 2026).

At the mechanistic level, these predictive mechanisms are thought to rely heavily on representational neural dynamics, specifically on the dynamic interplay between two classes of neural representations: conflict-expectancy representations, which encode the continuously updated probability of conflict derived from environmental regularities, and task representations, encoding the task-relevant (target) and task-irrelevant (distractor) information required for task execution (e.g., Badre et al., 2021; Cellier et al., 2022; Jiang et al., 2018). The core hypothesis, more or less explicitly postulated across theoretical accounts of cognitive stability (e.g., Braver, 2012; Egner, 2023; Miller & Cohen, 2001), is that predictive stability mechanisms rely on the encoding of conflict-expectancy representations, and that these, in turn, modulate the encoding strength of task representations directionally —strengthening the relevant task representation and weakening the irrelevant one— thereby shielding the task focus from conflict before it occurs. Yet, despite the theoretical centrality of this mechanism, its representational implementation has remained largely underspecified. This is partly because most prior work has focused on downstream conflict outcomes rather than upstream predictive dynamics. Moreover, even when MVPA analytical tools have been employed, prior work has provided neither a direct test of how expectations modulate task representations trial-by-trial, nor a fully multidimensional characterization of these representational dynamics that jointly leverages their spatial, temporal, and spectral patterns.

In the present study, we addressed these gaps by leveraging a perifoveal spatial Stroop paradigm in which global (LWPC) and item-specific (ISPC) conflict probabilities were concurrently manipulated to engage the two predictive stability modes at the same time. We modeled the trial-by-trial dynamics of both conflict-expectancy types with an ideal Bayesian observer, and applied Representational Similarity Analysis —complemented by convergent ridge-regression decoding— within an integrated spatio-temporo-spectral searchlight framework, allowing us to jointly track the representational dynamics of predictive stability across the spatial, temporal, and spectral domains with high resolution. Critically, our RSA approach was designed to identify not only whether task representations are modulated by conflict expectations, but also the direction of this modulation —a level of specificity that goes beyond the descriptive quantification of encoding-strength differences employed so far. Our findings reveal that predictive stability operates through the anticipatory and sustained encoding of conflict-expectancy representations, which directionally modulate task representations along dissociable dynamics for its proactive and predictive reactive forms.

Before detailing the functional architecture of our findings, their interpretation must be constrained by the physiological and inferential properties of non-invasive electrophysiology. First, in the temporal domain, reported peak coordinates index maximal observed representational encoding strength rather than isolated, punctate events; statistical inference applies to the temporal cluster as a whole. Second, in the spatial domain, sensor-level topographies are treated as descriptive operationalizations of multivariate scalp activity patterns and are not interpreted as precise cortical localizations. Third, main effects and interaction terms capture complementary aspects of the same representations: the former index where Task information is most strongly encoded on average across trials, the latter where its encoding strength scales with conflict expectations. Their patterns therefore need not coincide, as high conflict expectations may strengthen the Target (and weaken the Distractor) by modulating its encoding in spatio-temporo-spectral patterns other than those dominating its average encoding. Fourth, in the spectral domain, mapping representational dynamics onto canonical frequency bands provides a physiologically motivated interpretive framework (Cavanagh & Frank, 2014; Jensen & Mazaheri, 2010; Spitzer & Haegens, 2017), grounding abstract representational geometries into potentially related established neurocomputational mechanisms. Crucially, our approach avoids the classical pitfall of collapsing across spatial, temporal and spectral dimensions, maximizing representational interpretability and directly revealing the coordinated multidimensional dynamics that implement cognitive stability. Crucially, to reduce reliance on post-hoc reverse inferences (i.e., inferring cognitive processes solely from non-specific univariate power shifts), our multivariate RSA directly interrogates the high-dimensional geometry of theoretically defined model RDMs, the informational content of each cluster is formally and mathematically determined by the model it reliably encodes (e.g., Target direction, Distractor location, or Bayesian conflict expectations).

In our multidimensional searchlight, the spectral dimension poses the highest interpretative challenge. Nevertheless, Theta and Alpha oscillations provide physiologically motivated reference points, given their well-established associations with cognitive control and attentional selection, respectively. Specifically, Theta rhythms (4–7 Hz) operate as a canonical oscillation for signaling conflict and cognitive control demands (Cavanagh & Frank, 2014; Cohen, 2014), while also supporting the relational integration and updating of contextual trial history across memory-guided networks (Fell & Axmacher, 2011; Klimesch, 1999; Sauseng et al., 2005). Concurrently, Alpha-band dynamics (8–12 Hz) serve as a primary instrument of attentional selection through selective functional inhibition, dynamically routing information flow by suppressing task-irrelevant visual channels and shielding ongoing task focus against interference —the “gating-by-inhibition” framework (Jensen & Mazaheri, 2010; Klimesch et al., 2007).

In contrast to the relatively well-characterized associations of Theta and Alpha, the Beta band (13–30 Hz) encompasses a substantially more multifaceted functional repertoire across the neocortex. Its interpretation thus requires a more specific conceptual framework. Rather than reflecting a single, monolithic process, Beta oscillations serve as a versatile electrophysiological infrastructure for the endogenous implementation of top-down constraints, state representations, and neural gain control (Bastos et al., 2020; Miller et al., 2018; Spitzer & Haegens, 2017). At the computational level, this translates into a single core operation: the active stabilization (or “inhibitory clamping”) of a neural state to prevent unauthorized state transitions. Crucially, the specific phenotypic manifestation of this putative clamping mechanism is not unconstrained, but emerges from the principled convergence of three analytically distinct constraints: (1) Informational Content (defined by the model RDM), (2) Carrier Frequency (indexed by sub-band tuning), and (3) Spatio-Temporal Stage (defined by the relevant trial window and scalp topography).

Under this unified computational principle, and in line with converging theoretical and neurophysiological evidence suggesting a functional gradient across the beta spectrum, we propose an interpretive framework in which neocortical networks leverage a functional division of labor between two distinct beta sub-regimes:

1. Upper-Beta Rhythms (∼20–30 Hz) in Action Selection and Executive Control: Oscillations in this upper range —classically linked in intracranial and source-localized studies to fronto-striatal and premotor circuits— have been consistently shown to mainly operate over rapid temporal windows dedicated to action-oriented operations, including:

(i) *Action selection and S–R translation*: the active encoding, selection of goal-directed task sets and rules, and their translation into specific motor outputs (e.g., Buschman et al., 2012; Herding et al., 2016, 2017);
(ii) *Prefrontal executive braking*: The deployment of top-down inhibitory clamping and braking signals to suppress intrusive representations and resolve conflict prior to motor execution (i.e., during response inhibition and conflict resolution; e.g., Hwang et al., 2014; Wessel & Aron, 2017);
(iii) *Executive status quo and delay maintenance:* The active holding of abstract task sets and structured probability landscapes across inter-trial delays within protected prefrontal spectral regimes (Engel & Fries, 2010; Lundqvist et al., 2016, 2018; Miller et al., 2018).
2. Lower-Beta Rhythms (∼12–20 Hz) in Contextual Priors and Sensory Scaffolding: Conversely, slower beta oscillations —predominantly documented across sensory and associative temporo-parietal cortices— operate over more extended temporal windows suited for continuous state tracking and representational routing, supporting:

(i) *Endogenous sensory reactivation:* The reactivation, maintenance, and gain control of internal sensory templates and perceptual representations (e.g., Hanslmayr et al., 2012; Gelastopoulos et al., 2019; Spitzer & Haegens, 2017);
(ii) *Bayesian prior tracking and updating:* The endogenous tracking, trial-by-trial updating, and pre-stimulus reinstatement of higher-order statistical regularities and contextual priors (Chao et al., 2022; Kopell et al., 2011; Tavano et al., 2019);
(iii) *Dynamic representational routing:* The calibration of top-down gain and inter-areal communication, dynamically biasing downstream sensory and motor channels (e.g., Bastos et al., 2020; van Ede et al., 2018).

We adopt this principled, tripartite-constrained framework as an interpretive lens to organize our beta-related findings. Rather than forcing a rigid functional dichotomy, this taxonomy captures a continuous gradient from slow contextual integration to fast executive routing. Critically, it offers a mechanistically informed rationale for our observed stage-dependent representational modulations, characterizing how the brain systematically channels specific computational contents (model RDMs) into dedicated oscillatory regimes (Upper vs. Lower Beta) across the trial cycle (spatio-temporal stage), thereby orchestrating both proactive and predictive reactive adaptation. While specific attributions within this framework remain provisional and await direct empirical verification, its adoption provides a principled and parsimonious lens through which to interpret the diverse beta-related effects reported below.

In what follows, we discuss these findings in turn (see Table 1 for a summary of all effects and their proposed functional roles): first, how task representations themselves are encoded (Section 4.1); second, how conflict-expectancy representations are encoded and updated across the trial cycle (Section 4.2); and finally, how these expectancies directionally modulate task representations through proactive (Section 4.3) and predictive reactive (Section 4.4) mechanisms. Unless otherwise noted, all effects reported below survived cluster-based correction, were supported by convergent evidence from ridge-regression decoding, and met the selection criteria outlined in Section 3.

### 4.1. Task representations

Cognitive stability fundamentally operates by acting on the representations that encode the information required for task execution —the task-relevant representation (Target) to be prioritized, and the task-irrelevant one (Distractor) to be shielded against (e.g., Egner, 2023; Miller & Cohen, 2001). These task representations are the substrates on which any form of stability, especially predictive stability, is postulated to act. Establishing whether they are reliably encoded in the neural signal is therefore a necessary starting point, providing the essential empirical foundation for subsequently examining whether and how their encoding strength is directionally modulated by conflict expectations.

#### 4.1.1. Target representation

The Target representation was reliably encoded in a right fronto-central Beta3 (i.e., 25–30 Hz) pattern, peaking at ∼420 ms after stimulus onset. While operationally modeled via the arrow direction (the trial-varying relevant input), theoretically the Target reflects an integrated task-set representation that binds the sensory feature, the task rule, and the specific motor response (i.e., “execute action X when perceiving feature Y”; Kikumoto & Mayr, 2020). Such an integrated representation can be considered a computational prerequisite for applying the task-defined stimulus–response mapping and selecting the appropriate action. Consistent with this view, related stimulus–rule–response representations have been shown to be engaged during action selection and to predict trial-by-trial variability in response times (Kikumoto & Mayr, 2020; Takacs et al., 2020); similarly, the strength of task-set representations has been shown to predict performance and has been related to stimulus-and response-related coding (Hubbard et al., 2019). By revealing a reliable encoding of the Target representation, the present finding provides the task-relevant representational substrate that can subsequently be modulated by predictive stability mechanisms.

The emergence of this effect in the upper-beta range is consistent with proposals that upper-beta oscillations support action-oriented goal representations, directly encoding task rules and driving stimulus–response (S-R) translation (Buschman et al., 2012; Herding et al., 2016). Rather than reflecting mere passive maintenance, upper-beta patterns over fronto-central and premotor circuits are thought to mediate the active transition from task-relevant sensory coding to categorical response selection (Herding et al., 2017). Although the encoding pattern emerged already at 20 ms, the relatively late peak (≈420 ms) is temporally compatible with the post-stimulus window in which stimulus–response translation processes are typically executed and resolved (Takacs et al., 2020). Because the present effect was stimulus-locked, we interpret it as a late stimulus-driven stage in which stimulus, rule, and response information are integrated into an action-relevant code that drives subsequent response selection, becoming most strongly expressed when it must be deployed for response generation.

Notably, no significant Target encoding survived cluster-based correction in the response-locked analysis. This absence is unlikely to reflect a genuine disappearance of the representation. Rather, because the Target representation reflects the integrated S-R translation process —mapping the visual arrow direction to its corresponding motor effector— its formation is fundamentally stimulus-driven in its temporal profile. Once this S-R translation is resolved, the baseline encoding of the representation may become less prominent in the period immediately preceding execution. Crucially, however, what becomes response-anchored is not its encoding per se, but rather its modulation by conflict expectations. Consistent with this interpretation, as will emerge from the analyses of proactive (Section 4.3) and predictive reactive (Section 4.4) modulation, high conflict expectations trigger a robust endogenous reinstatement of the Target representation in the peri-response window. This indicates that the motor-anchored expression of the Target is not a tonic, demand-invariant encoding, but rather a strategic, demand-dependent amplification: the cognitive system dynamically re-instantiates the task-relevant S–R code precisely prior to execution only when competing distractor interference is anticipated.

#### 4.1.2. Distractor representation

Although cognitive stability fundamentally operates by prioritizing task-relevant representations, in a conflict paradigm like ours, it is highly probable that a task-irrelevant representation (the Distractor) is also co-encoded. Spatial position, in particular, is processed automatically and is virtually impossible to ignore entirely; the more automatic the Distractor, the stronger the resulting conflict. Of note, our experimental paradigm was specifically designed to elicit strong spatial conflict. In line with this, our results reveal robust Distractor representations in both stimulus- and response-locked data.

In the stimulus-locked data, the Distractor was encoded in a right (pre)frontal and central pattern within the lower-beta range (Beta 1-2; 13–24 Hz). The significant encoding peaked early (∼200 ms) but was also present in later time windows throughout the entire epoch. The early peak —occurring well before the 420-ms peak of the Target— is consistent with the rapid and automatic extraction of spatial location, demonstrating that irrelevant positional information is rapidly processed by early visuospatial systems and is notoriously difficult to fully suppress even when task-irrelevant (Cespón et al., 2016; Martín-Arévalo et al., 2015). Crucially, this temporal dissociation provides neural evidence that processing stimulus position is more automatic than processing stimulus direction, an assumption that we previously posited, but that had yet to assess empirically at the multivariate neural level (Viviani et al., 2024b; 2024c). Moreover, the temporal dynamics of the Distractor encoding aligns with the role of lower-beta rhythms in maintaining endogenous sensory traces and content representations (Hanslmayr et al., 2012; Spitzer & Haegens, 2017), as well as with recent multivariate evidence showing that task-irrelevant visual representations can remain decodable for up to approximately 1 second (Noah et al., 2023). Thus, this persistent distractor encoding may provide a substrate for continued conflict, while the later peak of the Target representation encoding described above may aid the resolution of this competition just prior to response execution. This conflict is likely further exacerbated by the topographical overlap between the two representations: both the Target and the Distractor are encoded by neural patterns with substantial overlap over right-lateralized frontal scalp sites, possibly related to underlying monitoring regions (Capizzi et al., 2016). When competing task-relevant and task-irrelevant features share overlapping spatial neural patterns, the potential for cross-talk and conflict increases substantially (Salzer et al., 2019).

To avoid overwriting within this shared topographical space, the system appears to rely on spectral dissociation. The task-irrelevant Distractor was encoded by lower-beta bands, consistent with the endogenous reactivation and maintenance of sensory and stimulus-specific content regardless of task-relevance (Hanslmayr et al., 2012, van Ede et al., 2018; see also Michelmann et al., 2022). The Target, in turn, appears to be expressed in upper-beta oscillations —as discussed above, action-oriented choices, task rules, and stimulus–response translation (Buschman et al., 2012; Herding et al., 2016, 2017). Whether this spectral separation reflects an active shielding mechanism or an emergent property of the distinct computational roles of the two representations remains an open question; regardless, it is compatible with the concept of frequency-specific routing and oscillatory multiplexing, whereby distinct frequencies support the routing of multiple information streams through partially overlapping neural substrates to prevent structural cross-talk (Akam & Kullmann, 2014).

Crucially, realigning the data to the response revealed a dynamic computational transition in how the Distractor is encoded as the trial unfolds. In the response-locked data, the Distractor was encoded in a circumscribed pre-response window (peaking at ∼-400 ms) over mid-central electrodes, within a higher frequency range (Beta2-3; 21–30 Hz). Importantly, because our multivariate models decode the specific spatial position rather than generic, non-specific motor preparation, this pattern reflects content-bearing information entering motor-related circuits. This topographical and spectral shift suggests that, while the stimulus-locked representation may reflect the persistent, stimulus-driven encoding of the irrelevant spatial feature and its automatic response mapping, the peri-response representation likely captures the critical moment when the Distractor propagates into the motor selection system. Observing the distractor encoded in central sensorimotor patterns just prior to response execution might reflect the critical transition where the irrelevant spatial code competes for motor output —a dynamic classically indexed by early incorrect motor activation in Lateralized Readiness Potential studies of spatial conflict (Valle-Inclán, 1996) and corroborated by electromyographic evidence of sub-threshold muscle activation preceding correct action execution (Burle et al., 2002). Furthermore, by shifting into the Beta2-3 range, the peri-response Distractor converges onto the spectral range in which the Target was encoded in the stimulus-locked data (Beta3). This suggests that, as the system approaches execution, the spatial distractor may transition into the same spectral range and fronto-central motor circuits, as goal-directed action plans do (Herding et al., 2016), potentially competing for response selection (a hypothesis that future work should test formally against trial congruency).

In summary, these results demonstrate that the brain reliably encodes both the task-relevant Target and the to-be-ignored Distractor as distinct multivariate patterns. The topographical overlap and peri-response spectral convergence of these two representations identify a candidate neural arena in which representational conflict may occur. Crucially, these task representations provide the representational substrate upon which predictive stability mechanisms are hypothesized to act, and whose modulation by conflict expectations is examined in the following sections.

### 4.2. Conflict-expectancy representations

To anticipate conflict before it fully emerges, predictive stability depends critically on representations of conflict-expectancy —internal, continuously updated estimates of the probability that conflict will occur, derived from environmental regularities (Egner, 2023; Jiang et al., 2014). To unveil the upstream mechanisms of predictive stability, two conditions must be met: first, conflict expectations must be encoded as distinct neural representations, defined by specific spatiotemporal and spectral patterns; second, these representations must be dissociable from the task representations themselves and contain sufficient information to drive the anticipatory modulation of task codes.

#### 4.2.1. Global conflict-expectancy (LWPC) representation

Proactive stability is theorized to rely on the sustained upregulation of task focus driven by high global conflict expectations (i.e., LWPC; Braver, 2012). Consistent with this framework, our analyses revealed that the continuously updated, trial-by-trial representation of global conflict expectancy (LWPC) is consistently encoded across both stimulus- and response-locked data.

LWPC was encoded already at stimulus onset and in a sustained manner in both stimulus- and response-locked data, with both representations peaking early (at ∼240 ms post-stimulus and ∼220 ms prior to the response, respectively). Crucially, its early onset and sustained expression extending towards the response, rather than being a transient reaction to stimulus onset, provide a candidate neural substrate for the implementation of proactive stability, in line with the DMC model (Braver, 2012). By being persistent, global conflict expectations can indeed function as anticipatory internal estimates that are accessed and utilized proactively to prepare the system (Egner, 2023), rather than acting as transient features triggered exclusively by stimulus identification. Importantly, because our trial-level LWPC predictor was derived from a dynamic Bayesian observer (HGF), its sustained neural encoding is not readily explained by simple time-on-task effects or linear drift alone, but is consistent with the active maintenance of a mathematically modeled belief state. Notably, the fact that LWPC encoding strength peaks well before the Target representation (which peaks at ∼420 ms) provides the necessary functional window for these global expectations to preemptively modulate the strength of task-relevant codes.

The functional nature of this proactive representation is further characterized by the specific spatial and spectral patterns encoding it. In the stimulus-locked data, LWPC was primarily encoded in the Theta band (4–7 Hz) across right (pre)frontal and left posterior spatial patterns. While midfrontal Theta has classically been characterized as a transient reactive alarm signaling the detection of acute conflict or errors (Cavanagh & Frank, 2014), growing evidence indicates that frontal Theta dynamics also track continuous fluctuations in sustained contextual control demands (Cavanagh & Shackman, 2015) and conflict expectation (Khan et al., 2025; Martinez-Molina et al., 2024). This is functionally consistent with the computational requirement of proactive stability: to track the occurrence of conflictual events as they unfold in order to maintain and update an internal representation of global conflict probability (LWPC). Thus, the frontal Theta pattern may reflect a representational structure that encodes this conflict evidence, providing the input necessary for the trial-by-trial Bayesian updating of global expectations. Concurrently, the LWPC representation was encoded by posterior Theta oscillations, which have been implicated in the maintenance and integration of trial-history regularities and working memory updating (Klimesch, 1999; Sauseng et al., 2005; see also Meng et al., 2021, for causal evidence linking left parietal Theta dynamics to associative and relational encoding). This functional interpretation aligns with the computational architecture of our paradigm, which requires the system to integrate the congruency of the current event with the mnemonic trace of recent history to update its internal model of conflict probability. Rather than isolated regional activity, the co-occurrence of these fronto-posterior Theta patterns is consistent with an integrated network supporting proactive state monitoring: while frontal circuits register ongoing control demands, posterior nodes maintain the historical evidence required to sustain a global, Bayesian model of conflict likelihood across consecutive trials (Fell & Axmacher, 2011; Jiang et al., 2015).

When locked to the response, the LWPC representation maintained its core Theta signature but became more widespread, extending to bilateral and central electrodes, while a prominent Alpha-band component also emerged. This spatial expansion is consistent with the broader engagement of core task-set maintenance and top-down control networks as proactive stability is implemented for action (e.g., Dosenbach et al., 2008; Hubbard et al., 2019). The preservation of the Theta pattern suggests that the same representational content is being tracked. Additionally, the concurrent emergence of an Alpha-band component may indicate a functional transition from representing an abstract expectancy to its operational implementation. Within the gating-by-inhibition framework (Jensen & Mazaheri, 2010), Alpha oscillations have been proposed to regulate the processing of task-irrelevant information by routing information flow away from task-irrelevant regions via selective functional inhibition. Accordingly, the emergence of Alpha within the response-locked LWPC pattern may reflect that, as the system approaches response execution, LWPC engages not only mnemonic representation (encoded in Theta), but also inhibitory gating signals (encoded in Alpha). This would effectively allow the LWPC representation to function as a proactive attentional filter, gaining the mechanistic capacity to bias sensorimotor processing by filtering task-irrelevant information at the point of action.

Taken together, the consistency of the LWPC representation across both stimulus and response levels —and its apparent unreliance from purely perceptual or motor dynamics— supports the view that LWPC is encoded as a sustained, content-specific neural representation that spans the full trial cycle, from conflict evidence integration to actionable control signaling to proactively enhance goal-directed task focus.

#### 4.2.2. Anticipatory trial-by-trial dynamics of LWPC representation

Beyond the sustained encoding observed during stimulus and response processing, our results uncover a dynamic, anticipatory trial-by-trial cycle of LWPC representation that appears to bridge successive trials. This cycle unfolds through three temporally distinct phases: late (post-response) event-related updating, sustained carry-over across the inter-trial interval (ITI), and pre-stimulus reactivation. Such a temporal dynamic provides a candidate mechanistic account of the representational dynamics supporting proactive stability, shifting the focus from static block-level states (e.g., Freund et al., 2021b; Gheza et al., 2026) to a continuous, trial-by-trial reconfiguration of the neural code.

The representational sequence begins late in the stimulus-locked epoch of trial *N* (∼800 ms post-stimulus, after response execution), where the LWPC representation is encoded in the Beta 1 band (13–18 Hz) within right temporal and mid-fronto-centro-posterior neural patterns. This late timing aligns with the possibility that the current trial’s congruency is incorporated into the history-dependent internal representation of global conflict probability (Egner, 2023; Jiang et al., 2014). Beta1 neural patterns are ideally suited for this role, as lower-beta rhythms have been implicated in the endogenous tracking, updating, and encoding of contextual regularities and statistical priors (Chao et al., 2022; Kopell et al., 2011; Tavano et al., 2019). This interpretation is reinforced by the pre-stimulus results of the following trial (*N+1*), which display topographical continuity with this stimulus-locked late cluster, suggesting a representational bridge between the two trials rather than two independent encoding events, and supporting the role of this late cluster as the starting point of a broader history-dependent carry-over mechanism.

Subsequently, during the ITI, the LWPC representation persists into the first pre-stimulus cluster of trial *N+1* (∼ 680 ms before stimulus onset), maintaining its right antero-temporal topography but shifting to an upper-beta range (Beta 2-3; 19–30 Hz). This pattern is consistent with the maintenance or reactivation of an abstract task-related representation within a protected, high-frequency oscillatory state during stimulus-free delays (Engel & Fries, 2010; Miller et al., 2018; Spitzer & Haegens, 2017). This phase therefore appears to serve as a representational bridge across successive trials, maintaining the updated global conflict-expectancy representation in a robust format across the inter-trial interval. Notably, the spectral difference (from later, post-response history integration in Beta1 to ITI persistence in Beta2-3) is consistent with a functional transition: while Beta 1 is optimally suited for computing and updating relational content tied to recent sensory evidence (Chao et al., 2022; Kopell et al., 2011; Tavano et al., 2019), higher-frequency beta rhythms (Beta 2-3) are typically recruited across frontoparietal networks to preserve the cognitive ‘status quo’ and shield internal control states against spontaneous decay during empty delay periods (Engel & Fries, 2010; Lundqvist et al., 2016; see also Miller et al., 2018 and Spitzer & Haegens, 2017). Finally, as the onset of trial *N+1* approaches (∼ -420 ms), the LWPC representation reconfigures into medial fronto-central patterns and, again, lower-beta oscillations (Beta1; 13–18 Hz). This transition is consistent with the endogenous reactivation and reinstatement of this global prior into an actionable, top-down predictive state (Bastos et al., 2020) that can subsequently be deployed to preemptively prioritize the Target representation (see Section 4.3.1).

Taken together, by being uncontaminated by stimulus-driven or response-related activity, these pre-stimulus findings provide multivariate evidence for anticipatory representational dynamics. This is theoretically significant: the encoding of conflict-expectancy representations before stimulus onset constitutes a necessary prerequisite for proactive stability, as the system can only preemptively reconfigure task codes if it has already encoded and made available the relevant expectancy information. Critically, this evidence challenges recent findings suggesting that conflict adaptation lacks anticipatory representational dynamics (e.g., Gheza et al., 2026). Our results demonstrate instead that global expectations are maintained as persistent neural representations throughout the trial cycle, including the pre-stimulus windows, and are kept available before the emergence of conflict. The overall observed spectral progression suggests that spectral neural dynamics scale with the current functional state of the representation. Rather than reflecting a static, block-level state that is suddenly switched on or off, this dynamic cycle suggests that the brain continuously tracks environmental regularities trial-by-trial: the global conflict-expectancy representations are updated after each trial based on experience, kept available across the inter-trial interval, and reinstated as actionable control information before each upcoming stimulus.

#### 4.2.3. Item-specific conflict-expectancy (ISPC) representation

Beyond global conflict expectations, the cognitive system also tracks conflict regularities tied to specific stimulus features. We propose that item-specific conflict expectancy —as operationalized here through the ISPC manipulation— drives a predictive reactive mode of stability: a mechanism that is triggered by stimulus identification but utilizes learned, item-specific conflict history to adaptively prepare the system before conflict fully emerges (Viviani et al., 2026). Therefore, if LWPC induces global-level stability adjustments, ISPC induces adjustments tailored to the identity of the current stimulus. Our analyses revealed that the neural patterns encoding trial-by-trial item-specific conflict expectations are consistently tracked across both stimulus- and response-locked data. Crucially, since all our multivariate models explicitly accounted for contingency —reflecting low-level stimulus-response associative learning (Schmidt, 2019)— these observed patterns represent a neural encoding of item-based conflict probability rather than simple S-R associations.

Temporally, the ISPC representation was encoded early after stimulus onset, peaking at ∼160 ms in the stimulus-locked data, and remained reliably represented through the peri-response window, with a second peak at ∼-260 ms relative to the response. The precocity of the initial stimulus-locked representation is consistent with the possibility that ISPC becomes neurally available shortly after stimulus identification, making the associated conflict expectation available at a timing compatible with what we term predictive reactive stability: reactive in that the representation is contingent on stimulus identification, yet predictive in that it may draw on learned item-specific expectations to establish an appropriate level of stability before conflict fully emerges. The persistent encoding of ISPC through the peri-response window further indicates that this representation remains reliably encoded during the critical stages of response selection, expressed in a form that could potentially support response-related processing. Taken together, these results suggest a two-step temporal profile of ISPC representation: an initial emergence shortly after stimulus identification, followed by sustained representation extending into the peri-response window.

Functionally, this two-step temporal sequence is characterized by a specific spatio-spectral encoding, showing a reconfiguration of the informational format of the item-specific conflict expectancy as the trial progresses. The early stimulus-locked ISPC patterns were predominantly expressed in the Alpha band over posterior electrodes. This posterior distribution is compatible with the involvement of visual regions in which environmental statistical regularities are encoded and modulate sensory representations (Ferrari et al., 2022; Kok et al., 2012; Summerfield & de Lange, 2014). The spectral encoding of this pattern is consistent with the role of Alpha oscillations in attentional gating, whereby task-relevant sensory processing is prioritized through the selective functional inhibition of task-irrelevant posterior areas (Jensen & Mazaheri, 2010). Therefore, given the ∼160 ms peak reported above, our results suggest that the ISPC representation likely encodes an inhibitory gating signal within the early stages of stimulus processing. Approaching the response, the ISPC representation maintains a posterior distribution but is expressed more medially and into lower-beta oscillations (Beta1). In line with our spectral framework, lower-beta rhythms have been associated with the endogenous retrieval, reinstatement, and maintenance of contextual priors and state representations (Bastos et al., 2020; Chao et al., 2022; Spitzer & Haegens, 2017). We thus tentatively propose that this spatio-spectral reconfiguration might reflect an algorithmic hand-off: the item-specific expectation is initially expressed in a form compatible with early sensory gating (Alpha) and subsequently reinstated in lower-beta as an endogenous prior representation, potentially contributing to conflict anticipation prior to response execution.

Overall, these spatio-spectral results point to a fundamental computational specialization between global (LWPC) and item-specific (ISPC) conflict-expectancy representations, uncovering a multi-level architecture for predictive stability. Spectrally, LWPC relies primarily on Theta oscillations, associated with sustained cognitive control monitoring and the integration of contextual trial history (Cavanagh & Shackman, 2015; Jiang et al., 2015; Sauseng et al., 2005), consistent with the need to maintain a macro-level, context-wide control policy; a supplementary Alpha component emerges only response-locked, when the maintained expectancy is translated into inhibitory gating for action. ISPC, by contrast, relies on initial Alpha oscillations, whose role in the selective gating of sensory processing (Jensen & Mazaheri, 2010) fits the item-specific nature of this expectation: ISPC likely encodes information used to selectively modulate the visuo-spatial processing of the specific item to which the expectation is tied, before being reinstated in lower-beta prior channels to inform decision circuits (Bastos et al., 2020; Spitzer & Haegens, 2017). Topographically, while LWPC was encoded by distributed fronto-posterior patterns characteristic of large-scale cognitive control networks (Dosenbach et al., 2008), ISPC remained confined to the posterior sensory-related patterns, reflecting localized statistical representations (Ferrari et al., 2022; Kok et al., 2012; Summerfield & de Lange, 2014).

Crucially, this dissociation provides a potential neurocomputational basis for distinguishing dual predictive stability modes: rather than reflecting a uniform expectancy signal, the results suggest a functional segregation between global proactive readiness (fronto-posterior Theta) and stimulus-triggered predictive adaptations (posterior Alpha/Beta1). Notably, Alpha oscillations contribute to both modes but in temporally distinct roles: as an early sensory gating signal associated with item-specific predictive reactive stability, and as a late, action-oriented gating component of global proactive stability.

#### 4.2.4. Anticipatory trial-by-trial dynamics of ISPC representation

Consistent with the anticipatory nature of predictive reactive stability, we further tested whether ISPC is also encoded in the pre-stimulus window as a signature of trial-by-trial anticipatory dynamics. Notably, unlike LWPC, no reliable ISPC pattern emerged in the late stimulus-locked window. This dissociation is likely explained by the specific statistical model tested in the stimulus- and response-locked analyses, which included only the ISPC of the current trial as predictor (see Section 2.4.2). Any post-response updating process, however, would necessarily concern the entire item-specific conflict probability mapping (ISPC_total) that must be carried forward to the next trial: the item just encountered on trial *N* acts as evidence for updating the full set of item-specific conflict probabilities, not only the entry associated with the current item. Consistent with this reasoning, when testing a model that included ISPC_total as we did in the pre-stimulus analyses, the trial-by-trial anticipatory dynamics of ISPC emerged. Two pre-stimulus clusters revealed a two-phase cycle across the inter-trial interval: an initial carry-over of the updated ISPC_total representation, followed by its reactivation approaching the imminent stimulus.

The first pre-stimulus cluster emerged early in the ITI (peaking at around −920 ms before stimulus onset), and was encoded in the Beta 2-3 band (19–30 Hz) over left anterofrontal patterns. This early cluster likely reflects the maintenance of the ISPC_total representation across the ITI through the abstract, working-memory retention of the latent multi-item probability states to keep the updated item-probability mapping available for the upcoming trial. Crucially, because our RSA model explicitly evaluated the similarity structure of the four-position multinomial distribution, this pattern captures the retention of a structured conflict-expectancy landscape rather than a non-specific, scalar state of uncertainty or arousal. Under our unified oscillatory framework, this reflects the deployment of upper-beta inhibitory clamping to stabilize the multi-item representation (ISPC_total) against decay during the stimulus-free delay, ensuring that the latent statistical structure of the environment remains protected across trials (Lundqvist et al., 2016; Miller et al., 2018; Spitzer & Haegens, 2017). This prefrontal Beta 2-3 signature mirrors the one identified for the analogous carry-over phase of the LWPC cycle, pointing to a shared upper-beta signature for the endogenous maintenance and protection of abstract task-set rules and conflict-expectancy representations.

The second cluster emerged later, closer to stimulus onset (peaking at around −220 ms), and displayed a widespread bilateral fronto-centro-posterior distribution and a mixed Theta-Alpha spectral profile. This peri-stimulus cluster likely reflects the reactivation of the ISPC_total representation in preparation for the upcoming trial. The fronto-central component resembles the analogous reactivation phase of the LWPC cycle, suggesting that both representations may become available for control at this stage (Bastos et al., 2020; Cavanagh & Shackman, 2015; Cavanagh & Frank, 2014). Unlike LWPC, however, ISPC_total reactivation extends to posterior patterns, potentially reflecting the greater representational complexity of ISPC_total: the reactivation appears to involve posterior visuo-spatial patterns associated with learned statistical regularities of the four possible positions (Ferrari et al., 2022; Kok et al., 2012; Summerfield & de Lange, 2014). Therefore, the initially abstract retention of the multi-item probability states encoded by earlier antero-frontal patterns is then translated into posterior visuo-spatial filters as the stimulus approaches, preparing the system for the specific spatial regularities of the four possible positions. Spectrally, the engagement of Theta oscillations is consistent with evidence that fronto-parietal Theta rhythms organize, bind, and maintain multiple distinct informational items within working memory (Klimesch, 1999; Lisman & Jensen, 2013; Sauseng et al., 2010, Sauseng & Liesefeld, 2020) —a computational requirement precisely tailored to ISPC_total, which demands the concurrent holding of a four-item probability distribution. Concurrently, the Alpha pattern is consistent with the attentional gating function established in the earlier stimulus-locked ISPC analysis (Jensen & Mazaheri, 2010).

This distinction resolves the apparent paradox of observing pre-stimulus representations for an inherently reactive control mode. We propose a two-tier architecture of predictive reactive stability that dissociates the maintenance of the complete item-probability landscape store in WM (proactive, pre-stimulus tier) from its item-specific deployment/reinstatement upon stimulus identification (reactive, post-stimulus tier). Crucially, this pre-stimulus encoding does not constitute proactive task preparation per se, as the upcoming stimulus identity remains unknown and the system cannot yet prioritize a specific directional response or filter a specific location. Rather, it serves as the essential predictive scaffolding: upon stimulus onset, the identification of a specific feature acts as a retrieval cue that immediately collapses this multi-item probability landscape into an item-specific control signal (ISPC_weighted), thereby triggering rapid, anticipatory task reconfigurations before conflict culminates.

Taken together, these findings provide the first multivariate evidence that ISPC, like LWPC, is also predictive rather than purely reactive: item-specific conflict expectancies are maintained dynamically across the inter-trial interval and reactivated in preparation for the next trial, rather than being confined to the current stimulus-response cycle. Together with the LWPC anticipatory dynamics described above, these dynamics point to a system in which multiple levels of conflict expectancy are continuously encoded, updated, and reinstated to preemptively prepare the brain for the upcoming trial. Having established that these two expectancy representations are reliably encoded and computationally distinct, the critical question becomes how these internal belief states are used upstream to directionally modulate task-relevant and task-irrelevant representations. We address this question in Sections 4.3 and 4.4.

### 4.3. Proactive modulation of task representations

Proactive stability is hypothesized to modulate task representations in an anticipatory and sustained manner, such that a high global expectation of conflict (low LWPC) dynamically strengthens the task-relevant representation (Target encoding) and weakens the task-irrelevant one (Distractor encoding) (Braver, 2012; Egner, 2023). Having established that LWPC is encoded as a specific representation reflecting the trial-by-trial global conflict expectancy and, therefore, the level of stability to engage (Sections 4.2.1-4.2.2), we examined whether and how this information is used to modulate task representations before conflict emerges, thereby characterizing the representational dynamics underlying proactive stability. Crucially, testing our hypothesis requires more than quantifying whether LWPC modulates the encoding strength of task representations: it also requires assessing whether such modulation follows the specific directions predicted by the theoretical framework (i.e., strengthening Target encoding and/or suppressing Distractor encoding). By explicitly capturing this direction, our results support the hypothesis, revealing multivariate neural patterns that proactively increase task focus and shield the system from distraction.

#### 4.3.1. Proactive stability increases Target encoding strength

The LWPC-driven modulation of the Target representation was characterized by an early-onset and sustained temporal profile. It emerged notably early (with two clusters peaking at ∼20 and ∼120 ms post-stimulus), persisted across the entire post-stimulus window (with a further cluster peaking at ∼480 ms), up to the peri-response period (cluster at ∼−140 ms), and was overall more strongly expressed in the stimulus-locked than in the response-locked analyses. This temporal dynamic suggests that the LWPC-driven Target strengthening is not a transient event triggered by the imminence of the response, but rather unfolds anticipatorily to accompany the selection process continuously, from stimulus onset up to its motor culmination, consistent with the anticipatory and sustained nature of proactive stability as formalized in the DMC model (Braver, 2012). Within this proactive profile, our findings revealed a coordinated two-tier oscillatory cascade operating across the trial cycle.

The first mechanism was expressed in the Beta 1-2 band, encompassing the early peaking stimulus-locked pattern (∼20 ms, over left centro-posterior electrodes) and a further peri-response one (∼−140 ms relative to the response, over medial posterior electrodes). Because the specific target feature (arrow direction) is unpredictable prior to display onset, the early stimulus-locked interaction cluster cannot reflect pre-stimulus encoding of the target identity itself. Rather, it likely captures the initial feedforward sweep of task-relevant sensory processing. Under predictive routing and biased-competition frameworks (Bastos et al., 2020), high global conflict expectations proactively establish an anticipatory top-down gain modulation. As incoming sensory inputs arrive, this baseline bias boosts the signal-to-noise ratio of task-relevant visual channels, resulting in an immediate amplification of Target representational discriminability from the earliest stages of cortical processing (Chao et al., 2022; Spitzer & Haegens, 2017). The re-emergence of this lower-beta pattern into the peri-response window further suggests that this task-relevant strengthening extends to the response preparation stage.

Alongside this task-relevant strengthening, the second mechanism, expressed in the Alpha band over left fronto-posterior patterns, was sustained throughout the stimulus-locked window (with peaks at ∼120 ms and ∼480 ms). In line with canonical frameworks of attentional gating and representational inhibition (Jensen & Mazaheri, 2010; Klimesch, 2012; Klimesch et al., 2007), this sustained Alpha pattern is consistent with active shielding of the prioritized target code against competing inputs. By maintaining continuous functional gating in favor of the Target representation, high proactive demand prevents representational interference. Notably, no analogous Alpha component was reliably encoded in the response-locked analysis, suggesting that this gating operates primarily as an upstream filter sustained across stimulus processing rather than a late-stage motor mechanism. However, null results should be interpreted cautiously.

Together, these complementary mechanisms show a structured spectro-temporal profile: an initial lower-beta predictive routing signal (∼20 ms) followed by sustained Alpha representational gating (∼120–480 ms) to shield target processing, and a pre-response lower-beta bias (∼−140 ms) to guide execution. Of note, all the stimulus-locked patterns supporting the LWPC-driven modulation of the Target were left-lateralized or medial over the scalp, a topographical signature that distinguishes them from the right fronto-central patterns encoding the Target representation itself (Section 4.1.1). This dissociation is consistent with the possibility that these patterns reflect a substrate expressing the LWPC-driven strengthening of Target encoding, rather than the encoding of the Target itself. This left-lateralized topography also partially overlaps with the posterior component of the LWPC representation (Section 4.2.1), which may suggest a functional continuity between encoding global conflict expectations and applying them to task-relevant codes.

#### 4.3.2. Proactive stability reduces Distractor encoding strength

While proactive stability primarily reinforces the task-relevant representation, it is complemented by a reduction of the Distractor representation encoding strength, which was temporally circumscribed and peri-response-locked.

The Distractor strength reduction unfolded in two complementary phases. First, a stimulus-locked pattern emerged late in the trial, approaching response execution (∼520 ms), in the Beta 3 band over right pre-frontal scalp patterns. This spatio-temporo-spectral neural code is consistent with the canonical electrophysiological signature of top-down inhibitory control and executive braking (Hwang et al., 2014; Swann et al., 2009; Wessel & Aron, 2017). Crucially, this upper-beta inhibitory signal is spectrally dissociated from the lower-beta range that encodes the Distractor representation itself (Section 4.1.2). This spectral dissociation suggests that right scalp prefrontal Beta3 activity does not reflect the passive decay of the distractor, but rather an active, top-down suppression of the interfering spatial representation before it reaches execution thresholds.

Subsequently, in the response-locked window, the modulation was expressed in the Theta and Alpha bands over left anterotemporal patterns. Because these are the same bands in which LWPC itself is encoded (Section 4.2.1), global expectancy information may interact with the Distractor representation through shared spectral patterns (Estefan et al., 2024; Jensen & Mazaheri, 2010). This spatio-temporo-spectral neural code is compatible with the possibility that global expectations actively suppress the distractor memory trace, attenuating the residual influence of the irrelevant spatial code on the final response selection cascade (Wöstmann et al., 2019).

This circumscribed temporal profile suggests a logic of computational economy underlying the LWPC-driven Distractor strength reduction: rather than being continuously downregulated throughout the post-stimulus window, the encoding strength of the Distractor is reduced specifically when the risk of interference with response selection is greatest. This reveals a fundamental computational asymmetry between task dimensions: the task-relevant Target requires continuous proactive amplification across the entire selection cascade (Section 4.3.1), whereas the task-irrelevant Distractor is managed through a proactively configured, late-stage brake that is triggered precisely when automatic spatial activations threaten to contaminate motor output (Hwang et al., 2014; Wessel & Aron, 2017).

Interestingly, our results complement recent findings suggesting that Distractor suppression during conflict adaptation lacks anticipatory representational states and operates purely reactively (Gheza et al., 2026). Here, we show that the peri-response reduction in Distractor encoding strength is proactively driven by LWPC, which is itself encoded as an anticipatory representation (Sections 4.2.1-2; see also Kelber et al., 2026, for computational evidence). Our findings thus reconcile these views: Distractor suppression emerges late in the trial sequence, which may explain why it appears reactive in coarse measurements, but its magnitude is governed by an upstream, proactive belief state that sets the gain of the putatively prefrontal inhibitory brake.

### 4.4. Predictive reactive modulation of task representations

Predictive reactive stability here is proposed to be a stimulus-triggered mechanism that leverages item-specific conflict expectancy (ISPC) to modulate task representations (Viviani et al., 2026), whereby the identification of a specific item associated with high conflict expectancy (low ISPC) should dynamically strengthen the Target encoding and reduce the Distractor encoding, through a representational dynamic triggered by stimulus identification that is distinct from the sustained proactive modulation. Having identified that ISPC is encoded as an anticipatory representation that exploits learned expectations while being triggered by stimulus identification (Sections 4.2.3-4), we here examined whether and in which direction this information is used to modulate task representations, in order to characterize the representational dynamics of the predictive reactive mode of stability. Our results support the hypothesis, revealing that ISPC directionally modulates task representations, with a peri-response-centred temporal signature that clearly distinguishes it from the earlier-onset and sustained proactive modulation examined above. These findings provide evidence for a dynamic form of stability that leverages item-specific information to increase task focus and reduce distraction.

#### 4.4.1. Predictive reactive stability increases Target strength

The ISPC-driven modulation of the Target representation was expressed later than the LWPC-driven modulation. While proactive Target strengthening was engaged from the earliest feedforward stages of stimulus processing (Section 4.3.1), the ISPC-driven strengthening reached its first reliable peak in the stimulus-locked patterns only at ∼460 ms —consistent with the fact that item-specific expectancies are contingent on stimulus feature identification. Notably, the peak of this modulation surfaced considerably later than the peak of the ISPC representation itself (∼160 ms; Section 4.2.3). This suggests that, although the expectation is encoded early, its operational deployment onto the Target becomes maximally expressed later, precisely when the Target may be most needed for response generation. Consistent with this interpretation, ISPC-driven Target reinforcement was more strongly and extensively expressed in the response-locked analysis, which encompasses two sub-clusters near the response (-160 and -60 ms).

This temporal profile mirrors and inverts the Target × LWPC profile: whereas LWPC modulation was more strongly expressed in the stimulus-locked analysis, extending continuously from stimulus onset and progressively narrowing toward the response, ISPC-driven Target strengthening was concentrated in the peri-response window, with only a stimulus-locked precursor, and peaked before the response. This inverted distribution may reflect the functional distinction between proactive and predictive reactive stability: sustained biasing (proactive) vs stimulus-contingent later boosting (predictive reactive). Functionally, this ISPC-driven Target reinforcement appears to be supported by two distinct but complementary mechanisms.

The stimulus-locked modulation, peaking at ∼460 ms, was expressed in the Theta band over right centro-parietal patterns, consistent with Theta’s proposed role in conflict-triggered control and reconfiguration (Cavanagh & Frank, 2014; Cohen, 2014). Specifically, upon identifying an item associated with a high conflict expectation (low ISPC), this spatio-temporo-spectral pattern may signal increased response selection demands and boost the task-relevant rule code (Takacs et al., 2020), precisely when interference from the spatial distractor is greatest.

The response-locked modulation, which captures this demand-dependent endogenous reinstatement, was strongly expressed in the lower Beta range across two right-lateralized patterns: a fronto-temporal pattern (peaking at −160 ms) and a centro-parietal pattern (peaking at -60 ms). In line with our oscillatory framework, the transition into lower-beta patterns suggests that, before response execution, item-specific conflict expectations, functioning as higher-order priors, increase Target representation discriminability to impose a robust pre-response bias (Chao et al., 2022; Spitzer & Haegens, 2017). Consistent with this account, the topographical progression from right fronto-temporal to right centro-parietal patterns may thus reflect the operational deployment of this strengthened Target representation, transitioning from working-memory substrates (Miller et al., 2018) to stimulus–response translation circuits (Takács et al., 2020).

When viewed alongside the proactive results, these findings reveal a double dissociation between proactive and predictive reactive stability: temporally, proactive stability (Target × LWPC) operates upstream as an early-onset, sustained baseline modulation, whereas predictive reactive stability (Target × ISPC) acts downstream as stimulus-triggered, peri-response reinforcement. Topographically, this dissociation is particularly evident in how the task-relevant Target representation is prioritized: proactive Target reinforcement is orchestrated through left-lateralized fronto-posterior scalp regions (Section 4.3.1), whereas item-specific Target boosting relies on right-lateralized scalp centro-parietal and fronto-temporal patterns. This topographical segregation suggests that proactive and predictive reactive mechanisms recruit distinct cortical routing channels to selectively amplify task-relevant codes at different stages of the selection cascade.

#### 4.4.2. Predictive reactive stability reduces Distractor strength

Beyond increasing the encoding strength of the Target, ISPC exerted an even earlier and more direct modulation on the Distractor, reducing its encoding strength. This was characterized by a remarkably precocious temporal profile, emerging earlier than the corresponding Target reinforcement in both stimulus-locked (∼300 ms vs. 460 ms) and response-locked (∼−320 ms vs. −160 ms) patterns. This stimulus-locked Distractor reduction closely followed the peak of the Distractor encoding itself (∼200 ms; Section 4.1.2), suggesting that the modulation acts on the Distractor representation shortly after it is fully expressed, precisely when it is maximally intrusive.

This reveals a “Distractor-first” functional hierarchy within predictive reactive stability: when a stimulus associated with high conflict probability is identified, the cognitive system prioritizes the rapid dampening of the specific interfering feature (the spatial position) before mobilizing resources to boost the task-relevant target code. This stands in striking contrast to proactive stability, which follows a “Target-first” policy (upregulating the Target from the earliest stages of stimulus processing (Section 4.3.1), while holding distractor suppression until late decision stages at ∼520 ms). This ISPC-driven Distractor reduction unfolded through two temporally distinct functional phases.

The initial phase comprised two complementary parallel mechanisms captured in the two locking analyses. In the stimulus-locked window, reduced Distractor strength was expressed in the Beta 1 range, peaking at ∼300 ms over left parietal patterns. The spectral overlap between this modulation and the lower-beta range encoding the Distractor representation itself (Section 4.1.2) is consistent with a stage-dependent targeted spectral gating (Akam & Kullmann, 2014): predictive reactive stability would dampen the intrusive spatial trace through a direct representational impact, operating within the same oscillatory signal as the representation it modulates. This would effectively reduce Distractor encoding strength at the perceptual level as soon as the item is identified. Concurrently, a response-locked pattern emerged (peaking at ∼−320 ms) in the Theta and Alpha bands over right posterior patterns, consistent with the Theta-mediated retrieval of item-specific priors (Klimesch, 1999) to implement early Alpha-mediated posterior attentional gating before motor commitment (Jensen & Mazaheri, 2010).

The later peri-response mechanism was expressed in the Beta 3 band over left parietal patterns, peaking at ∼−100 ms. Remarkably, this late modulation mirrors the second stage of our spectral gating framework: just as the Distractor representation shifts into upper-beta frequencies (Beta2-3) near response execution to compete for motor selection (Section 4.1.2), predictive reactive control re-emerges in the Beta3 band to exert a final pre-response brake (Hwang et al., 2014; Wessel & Aron, 2017). By shifting from Beta1 (∼300 ms) to Beta3 (∼−100 ms), the control system matches the evolving spectral signature of the Distractor. Functionally, this late parietal Beta3 pattern reflects the final stage of localized inhibitory clamping, applying a targeted pre-response brake that silences the irrelevant spatial code precisely as it competes for access to motor execution channels. The consistent involvement of left parietal patterns across both latencies suggests topographic continuity between the two phases, reflecting a unified parietal control loop that tracks and suppresses the spatial distractor from its initial perceptual representation to its motor culmination.

Overall, ISPC-driven Distractor reduction was expressed over predominantly parietal patterns, consistent with the visuo-spatial processing of the Distractor feature in the paradigm employed here (i.e., the arrow position). Interestingly, this provides a further topographical and computational dissociation from proactive modulation of the Distractor (Section 4.3.2): LWPC engages a right frontal executive braking alongside left temporal suppression, whereas ISPC operates locally within posterior visuo-spatial parietal circuits via stage-dependent, targeted spectral gating. Together with the Target results, the ISPC-driven modulations of Distractor encoding suggest that predictive reactive stability is not merely a weaker or delayed copy of proactive stability, but an autonomous, highly specialized mode governed by a “Distractor-first” logic and implemented through frequency-matched representational suppression.

### 4.5. Proactive and predictive reactive stability: A multidimensional representational architecture for conflict anticipation/mitigation

Taken together, our findings uncover a highly dynamic and adaptable representational architecture for cognitive stability. Rather than relying on static, block-level control states, the brain dynamically adjusts the neural encoding strength of task representations on a trial-by-trial basis, scaling its modulations in proportion to continuously updated Bayesian conflict expectations — global for LWPC and item-specific for ISPC. Crucially, although both stability modes share the same normative objective —strengthening the task-relevant Target and weakening the task-irrelevant Distractor— our multidimensional searchlight reveals that they achieve this outcome through two computationally distinct policies:

1. Proactive Stability operates via a “Target-First / Global Upstream Bias”. Driven by sustained fronto-posterior Theta and pre-stimulus Beta priors (Section 4.2.1-2), proactive stability deploys an early lower-beta routing signal and sustained Alpha representational gating to continuously amplify the Target code across the entire selection cascade (Section 4.3.1). In contrast, Distractor suppression is withheld until late peri-response stages (∼520 ms), where an upper-beta prefrontal brake is recruited to prevent the spatial feature from invading motor channels (Section 4.3.2). This reflects a policy of global proactive shielding, where cognitive resources are invested primarily in maintaining and shielding the task-relevant focus.
2. Predictive Reactive Stability operates via a “Distractor-First / Local Spectral Gating”. Driven by the rapid collapse of the pre-stimulus item-specific conflict-expectancy landscape upon item identification (Section 4.2.4), predictive reactive control prioritizes the immediate neutralization of the intrusive feature (Section 4.4.2) before boosting the target (Section 4.4.1). It achieves this through stage-dependent targeted spectral gating within posterior circuits, directly suppressing the Distractor’s sensory codes and subsequently limiting its motor expression.

This dissociation provides novel insights beyond classical models of cognitive control, most notably the DMC framework (Braver, 2012). Classically, reactive control has been conceptualized almost exclusively as a late, corrective “just-in-time” braking mechanism engaged after conflict has already disrupted processing (Braver et al., 2007). Our findings provide evidence consistent with the existence of an intermediate predictive reactive mode: a mechanism that is triggered reactively by stimulus identification, but operates predictively by retrieving learned item-specific priors to reconfigure task codes before downstream corrective processes become necessary (Viviani et al., 2026). By uncovering these upstream representational trajectories, our results suggest that conflict adaptation in item-specific paradigms is not a passive, downstream outcome, but may reflect structured, expectancy-driven representational dynamics.

#### 4.5.1. Limitations and future directions

Several methodological and theoretical limitations of the present study warrant consideration.

First, as noted at the beginning of the Discussion, although our 3D searchlight framework integrates spatial, temporal, and spectral dimensions, scalp EEG inherently reflects mixed field potentials subject to volume conduction. Consequently, our topographical descriptors (e.g., “fronto-central”, “posterior”) refer to sensor-level representational patterns, not millimetric anatomical structures or deep subcortical generators (such as striatal nodes). Future studies combining high-density M/EEG with structural source modeling or concurrent fMRI will be essential for mapping these distinct oscillatory representational channels onto specific subcortical and laminar microcircuits.

Second, our findings are based on a perifoveal spatial Stroop paradigm, which is well suited to isolating automatic spatial interference. Establishing whether this dual-tier architecture generalizes to other cognitive domains (e.g., semantic Stroop, Flanker, or task-switching paradigms) is an important avenue for future research.

Finally, while the Hierarchical Gaussian Filter provides a principled, normative account of trial-by-trial statistical learning, exploring alternative computational learning models (e.g., reinforcement learning with dynamic learning rates) may offer complementary perspectives on how the brain learns and updates conflict expectations.

## 5. Conclusions

The present study provides the first comprehensive, multivariate characterization of the representational dynamics underlying predictive cognitive stability. By combining an integrated spatio-temporo-spectral searchlight RSA approach with trial-by-trial Bayesian modeling of conflict expectations, we directly tested and found support for the core theoretical hypothesis that predictive stability operates by dynamically and directionally modulating the neural encoding strength of task representations based on continuously updated conflict expectations.

These findings carry three implications for cognitive neuroscience. First, we bridge the long-standing gap between theoretical postulates of representational control and empirical observation. Stability is not merely indexed by reduced conflict or reaction-time adjustments downstream, but is implemented upstream through the active, directional modulation of task-relevant (Target enhancement) and task-irrelevant (Distractor weakening) neural patterns. Second, our findings are consistent with the view that conflict adaptation may rely on anticipatory representational dynamics. The human brain does not treat control demands as discrete, static blocks; rather, it maintains a continuous Bayesian scaffolding across the trial cycle, updating conflict expectations post-response, preserving them across inter-trial intervals, and reinstating them as actionable top-down priors before stimulus onset. Third, we formalize a critical distinction between proactive stability (a Target-first global bias induced by LWPC) and predictive reactive stability (a Distractor-first local policy induced by ISPC).

Lastly, from a methodological perspective, our multi-feature searchlight framework establishes an analytical template for interrogating the high-dimensional neural code of the human brain. By simultaneously tracking information across space, time, and canonical oscillatory rhythms, this approach opens new avenues for uncovering the underlying computational mechanisms that orchestrate goal-directed behavior across diverse domains of human cognition.

## Footnotes

1 While the DMC framework refers to these modes in terms of cognitive control, they can be directly applied here to the domain of stability.

